# The Winners Take It All? Global Evolutionary Success of H5Nx Reassortants in the 2020–2024 Panzootic

**DOI:** 10.1101/2025.07.19.665680

**Authors:** James Baxter, Jing Yang, William Harvey, Simon Dellicour, Anne Pohlmann, Mingxiao Ma, Jinhua Liu, Wenjun Liu, Yuhai Bi, Paul Digard, Martin Beer, Marion Koopmans, Samantha Lycett, Lu Lu

**Author notes:** These authors contributed equally to this work (James Baxter, Jing Yang).

## Abstract

Avian influenza viruses undergo frequent genetic reassortment, which can coincide with phenotypic changes in transmission, pathogenicity, and host species niche. Since 2020, clade 2.3.4.4b H5 high pathogenicity avian influenza viruses (HPAIVs) have driven a global panzootic, causing mass mortality in wild birds, poultry, and, for the first time, repeated spillover infections in a variety of mammalian species. This worldwide resurgence of H5 HPAIV has coincided with a dramatic increase in the number of circulating reassortant strains; however, the scale, impact and drivers of these reassortants remain unclear. Here, we combined statistical and phylodynamic modelling to reconstruct the global evolutionary dynamics of H5Nx viruses across four epizootic seasons (2020-2024). We identified 209 genetically distinct reassortants, stratified into three transmission categories based on their phylogenetic and epidemiological profiles. Accounting for sampling depth and HPAIV incidence, we estimated that reassortants emerged most frequently from Asia, but ‘major’ reassortants associated with increased host range, inter-seasonal persistence, and long-range dissemination, more frequently emerged from Europe. Altogether, reassortant emergence followed an episodic pattern in which most reassortants were transient, but 2% seeded large clusters of secondary reassortants soon after their own emergence. Statistical modelling revealed that reassortant success was strongly shaped by ecological factors, including sustained circulation in specific wild bird orders and detection across a wider range of host niches. Collectively, our findings uncover global reassortment dynamics in H5 HPAIVs and identify key virological and ecological drivers underpinning the emergence and spread of successful reassortants. These insights support the importance of enhanced surveillance to track evolution of H5 HPAIV and identify traits relevant for consideration in pandemic risk assessment.

## 1 Introduction

The Influenza A virus genome comprises eight single-stranded negative sense RNA segments, which can reassort when two or more strains coinfect the same cell [1]. Genetic reassortment can alter Influenza A virus evolutionary dynamics, if it leads to phenotypic ‘shifts’ that play a critical role in the emergence and cross-species transmission of novel viral strains [2]. Notably, reassortment of Influenza A viruses has been implicated in several past pandemics including 1918 H1N1, 1957 H2N2 and 1968 H3N2, where gene segments from avian influenza viruses combined with human-adapted viruses [3–5]. While inter-subtype reassortment between Influenza A viruses circulating in the human population is relatively infrequent, the coexistence of multiple Influenza A subtypes in the wild avian population coincides with a remarkably high rate of reassortment [6, 7].

H5 High Pathogenicity Avian Influenza Virus (HPAIV) was first isolated from a poultry outbreak in Aberdeenshire, Scotland in 1959 [8]; however, it was not until the H5N1 goose/Guangdong (Gs/Gd) lineage emerged in China in 1996 that sustained transmission in domestic poultry was established [9]. Since 2005, repeated spillovers from domestic poultry to wild Anseriformes (geese, ducks and swans) established a broad diversity of H5Nx lineages across Asia, Europe, and Africa [10–12]. In 2014, the emergence of a lineage with reduced virulence in wild Anseriformes (clade 2.3.4.4), led to the introduction of Gs/Gd HPAIV to North America for the first time, circulating until 2015 [13, 14]. From 2016 onwards, clade 2.3.4.4b H5N8 viruses originating from East Asia repeatedly caused outbreaks in wild Anseriformes, coinciding with a relative increased virulence for Anatidae spp and a heightened reassortment frequency compared with the 2014/2015 seasons and earlier European H5N1 outbreaks [15–17].

Since 2020, clade 2.3.4.4b H5N1 have surged to cause unprecedented outbreaks in wild-bird species worldwide, displacing contemporary H5N8 lineages, and showing remarkable persistence in wild birds alongside a broadened host range [18–21]. By 2024, descendants of the 2020 H5N1 lineage had caused severe mortality in wild birds and domestic poultry across Europe, Africa and latterly North America, following the reintroduction of H5N1 in 2021. Moreover, sustained avian-to-mammal transmission for the first time became apparent across multiple settings including fur farms [22], as well as wild marine and terrestrial mammals [23–27]. Circulation in mammals, notably cattle in the USA, has resulted in sporadic mammal-to-human transmissions, typically following prolonged exposure [28, 29].

The 2020-2024 panzootic was characterised by extensive genetic reassortment and circulation of reassortant viruses across a wide diversity of host species worldwide. The reassortment dynamics of clade 2.3.4.4b H5Nx viruses from Europe in 2020–2022 [30] and from North America [31, 32] have been reported; however, a systematic investigation of this complex evolutionary picture at the global scale has yet to be undertaken. In this study, we updated a previously described global reassortant classification system [16] to characterise H5 HPAIV reassortants of clade 2.3.4.4b that circulated between 2020 and 2024. We used phylogenetic approaches to ascertain spatio-temporal patterns of reassortant emergence, before combining phylodynamic and statistical modelling to elucidate drivers of reassortant persistence and dispersal. We show that a small minority (5/209) of reassortants dominated the 2020-2024 panzootic, and reveal how short inter-reassortment generation times led to a clustered, episodic pattern of reassortant emergence. Averaged across all reassortants, we find viral dispersal was maintained in Anseriformes spp, and potentially accelerated by circulation in Charadriiformes spp. Collectively, we discuss the role of virological and ecological factors on the formation of distinct reassortment patterns throughout the 2020-2024 panzootic.

## 2 Methods

### 2.1 Sequence Data Curation and Assembly

We assembled a dataset of all available HPAIV H5 clade 2.3.4.4b full genome sequences with at least 85.0% nucleotide coverage from the GISAID EpiFlu database collected between 1st Jan 2019 and 1st May 2024 (Downloaded 28th May 2024) [33]. These data comprised 9,935 Influenza A virus genomes, including viruses of the subtypes H5N8 (*n* = 1,587), H5N1 (*n* = 8,143), H5N5 (*n* = 94), H5N6 (*n* = 70) and other H5Nx (*n* = 41). 93.9% of these sequences were sampled from birds (*n* = 9,325), with the remainder sampled from mammals (*n* = 457) and the environment (*n* = 153). Environmental samples included those sourced from water, faeces and bird habitats (e.g. cages). Our data also included early sequences sampled from H5N1 infected cattle in the United States of America (*n* = 12, collected in March 2024). Altogether, most Influenza A virus genomes in our data were sampled from Europe (*n* = 4574), followed by North and Central America (*n* = 3311), Asia (*n* = 1566), and South America (*n* = 187). The majority (87.1%) of whole genomes were sampled between 2021 and 2023, with the number of isolates sampled from Asia and Europe typically peaking during northern-hemisphere winter (Figure S1) . Details of data and data providers are summarised and acknowledged (Table S1).

### 2.2 Reassortant Classification

We determined reassortment profiles for each influenza genome according to a previously described reassortant classification algorithm for global clade 2.3.4.4b H5 AIVs [16]. Briefly, we aligned the nucleotide sequences of each of the eight gene segments using MAFFT v7.511 [34]; excluded noncoding regions and removed insertions present in fewer than 10% of sequences. We defined clusters using nearest neighbour hierarchical clustering for all whole genomes in our dataset. Specifically, we clustered pairwise nucleotide distances, setting a 0.5% threshold for the longest three gene segments (PB2, PB1 and PA) and a 1% threshold for the remaining segments using “bioseq” package in R v4.1.2 [35]. We selected different thresholds to account for differences in segment length, resulting in monophyletic clusters distinguished by approximately 8-15 nucleotides (Figure S4). For each gene segment cluster, we assigned a number that corresponds to the ranked cluster size (1-n, largest to smallest), resulting in a unique 8-number barcode based on the cluster combination of eight genomic segments for each genome (e.g 21111111). For ease of reference, the label for each reassortant profile is given by the virus HN subtype, collection year of the first sampled genome, and the chronological order of discovery among all reassortant profiles identified within that year (e.g. 2020/H5N1/R1). Our reassortant profiles were broadly consistent with genotyping schemes established by the European Union Reference Laboratory for Avian Influenza and Newcastle Disease [36] and the United States Department of Agriculture [31] (Figure S3).

### 2.3 Time and location-stratified reassortant diversity

We investigated the diversity of H5Nx reassortants through time, stratified by their sampling continent. Using a sliding time window of one year, we calculated the proportional abundances of reassortants sampled in each affected continent, excluding Antarctica (Africa, Asia, Europe, North America and South America). From these proportional abundances, we calculated diversity as the Hill number of order 1 [37]. This measure accounts for both the number of reassortants present within a discrete place and time and the evenness of the spread or distribution of reassortants; it represents the effective number of reassortants present if they were evenly distributed and is equal to the natural exponent of Shannon entropy. To mitigate the assumption that all reassortant profiles are equally distinct, we additionally calculated a similarity-sensitive measure of diversity [38]. We calculated a similarity matrix from pairwise nucleotide distances between consensus sequences for each reassortant using an exponential transformation using the “rdiversity” package in R v4.1.2 [39]. To account for differences in sampling intensity between continents, we repeatedly and randomly down-sampled overrepresented continents to match the number of profiles present in the least well represented continent. For each metric, we quantified patterns over time using a univariate generalised additive model (GAM) (using “mgcv” v1.9-1 package in R v4.1.2) with a penalised cubic regression spline fitted to the midpoints of the sliding windows [40].

### 2.4 Phylodynamic analyses

To provide a contextual background for further analyses, we identified the 500 most genetically similar sequences for each gene segment of the earliest isolate of every reassortant profile using the Basic Local Alignment Search Tool (BLAST) [41]. We used the default parameters provided by the GISAID database and did not constrain our search results by host, location or subtype. We included only sequences sampled on or before the collection date of the reassortant isolate and included only one of any duplicated sequences based on the isolate names for each gene segment. We curated the accompanying data for the contextual sequences as described for our whole genome data.

To infer key evolutionary parameters of all reassortant profiles identified in this study, we first grouped genomes into reassortants that were found within a given continent (intracontinental) and reassortants found in more than one continent (intercontinental). We analysed intercontinental reassortants (*n* = 13) individually and analysed intracontinental reassortants (*n* = 196) together within each continent. For each dataset, we analysed each segment and NA serotype separately. We inferred maximum likelihood trees using IQTREE v2.1.3, assuming an HKY (Hasegawa Kishino Yano) substitution model with a four-category gamma distribution model for among-site rate heterogeneity [42–44], and confirmed the temporal signal of each alignment using the root-to-tip regression implemented in Tempest v1.5.3 [45].

To balance the distribution of sequence samples across each phylogeny, we sub-sampled each alignment. Specifically, we calculated pairwise Hamming distances (HD) and inferred sequence clusters with a maximum difference of 5 SNPs. We then selected one genome for each unique combination of location (defined as most refined administrative subdivision available), host order, reassortant, and HD cluster.

We estimated Bayesian time-resolved phylogenetic trees for each alignment using BEAST v1.10.4 coupled with the BEAGLE v3.1 library to enhance computational performance [46, 47]. We assumed a SRD06 nucleotide substitution model, in which codon positions 1+2 and 3 are partitioned and a separate HKY model with a four-category gamma distribution model for among-site rate heterogeneity, is fitted to each [42, 43, 48]. Additionally, we assumed an uncorrelated relaxed molecular clock, with evolutionary rates sampled from a lognormal distribution [49]. We specified a lognormal prior for mean evolutionary rate (*X* ∼ LogNormal(−8.537, 1.805)), a uniform prior for relative rates amongst partitions (*X* ∼ Uniform(0, 100)) and a nonparametric skygrid tree prior [50]. For each gene-segment alignment, we ran two independent Markov Chain Monte Carlo (MCMC) simulations, each comprising 2 × 10^8^ iterations with sampling every 20,000 iterations. We assessed convergence and satisfactory effective sample size (ESS >200) using Tracer 1.7.2 [51].

For each fitted phylogeny, we inferred the ancestral states for five key traits. First, we estimated the most recent common ancestor (MRCA) for every reassortant in each phylogeny, reconstructing the ancestral reassortant ‘state’ using a symmetric continuous-time Markov chain [52]. Next, we conducted further discrete trait analyses for influenza subtype, sampling continent, and sampled host, each time assuming an asymmetric continuous-time Markov chain. Finally, we modelled the fine-scaled continuous diffusion patterns for each reassortant using a gamma-distributed relaxed random walk model [53]. We established the geospatial location (latitude and longitude) of each isolate through extensive literature searches. Where the precise coordinates could not be uniquely identified, we used the centroid of the best-resolved (county/state/country) polygon. Where tip-locations were identical, we modelled a random jitter of 0.001. For all reassortants, we estimated the weighted diffusion coefficient, which approximates the area invaded by sampled lineages per unit of time [54, 55].

We combined these analyses to create a detailed phylodynamic profile for each reassortant. To disentangle reassortant-specific ancestral states and persistence times, we identified the most recent common ancestor for each reassortant across a 1000 tree subset of the posterior tree distribution. We recorded the posterior estimates for the node age, host identity, location and subtype and then calculated the evolutionary time each reassortant spent in each host-state, the total evolutionary ‘persistence’ time (time between the most recent sampling date and the estimated date for the MRCA), the frequency of host transitions, and the frequency of avian-to-mammal transitions. In the main text, we report the median values and summarise each posterior tree distribution with a maximum clade compatibility tree, estimated using TreeAnnotator (v1.10.4) [56].

To better understand underlying factors related to the dispersal dynamics of the five major H5Nx reassortants of clade 2.3.4.4b identified in this study, we extended our discrete trait analysis with a phylogeographic generalised linear model (PGLM) [57]. Specifically, we modelled the transitions between geographical regions adapted from the UN Geoscheme (Figure S26), as a function of eight geospatial predictors: i) the number of intersecting terrestrial flyways between regions (https://datazone.birdlife.org/about-our-science/flyways); ii) the median density of chickens (number of animals per km2) and iii) the median density of ducks, each summarised from Gridded Livestock of the World Database 4 across each region [58]; iv) the great-circle distance between the centroids of each region; v) the ratio between the coastline length (https://www.wri.org) and area (https://www.cia.gov/the-world-factbook), vi) the median of the mean annual temperature [59], vii) the proportion of the area of covered by wetlands as classified by the Global Lakes and Wetlands Database [60]; and viii) the number of whole genomes sampled. We standardised these continuous variables to a mean value of 0 and a standard deviation of 1. Prior to model fitting, we tested for multicollinearity between the variables (Figure S28). If the absolute Pearson correlation coefficient between each pair of predictors was greater than 0.7, these predictors were considered highly correlated and included in separate PGLMs to mitigate the influence on coefficient estimation. We pre-processed spatial data in R using packages sf v1.0-20 [61], terra v1.8-42 [62] and tidyterra v0.7.2 [63].

### 2.5 Statistical analyses

#### 2.5.1 Number of Reassortants

To understand global patterns of reassortant emergence, we fitted a hierarchical model to predict the number of novel reassortants, *y*, stratified by year-month, *i*, and continent, *j*. Our model consisted of three components, inspired by ecological abundance models [64, 65]. First, we hypothesised that reassortants could only emerge at a fraction of time points, perhaps due to epidemiological or ecological suitability. We assumed conditions at each time point were either permissible or not for reassortment, *z*_*ij*_ ∈ {0, 1}, with continent-specific probability, *θ*_*j*_ ∈ (0, 1). Next, we assumed that only a proportion of reassortants, *p*_*ij*_ ∈ (0, 1), are ever observed due to incomplete sampling. Finally, we assumed that a ‘true’ latent number of reassortants per month per continent, *N*_*ij*_, follows a Poisson distribution with continent-stratified rate *λ*_*ij*_.

We modelled additional linear predictors for each of *θ*_*j*_, *N*_*ij*_ and *y*_*ij*_. Specifically, we included continent-specific intercepts for all components, in addition to global coefficients for HPAIV incidence and the number of GISAID whole genomes for *y*_*ij*_ and *N*_*ij*_, respectively. We approximated HPAIV incidence as the number of (exposed) subunits from all biannual reports submitted to the World Organisation for Animal Health (WOAH) related to H5 HPAIV in wild birds, mammals and poultry between 1st January 2017 and 31st December 2024 (Accessed 2025-Feb-25) (Figure S1). We log-transformed continuous predictors prior to fitting the model. We fitted non-centred random intercepts for the year in which the reassortant was estimated to have emerged for *N*_*ij*_ and *y*_*ij*_. A full description of the statistical model is provided in the supplementary methods (Appendix A.1).

#### 2.5.2 Reassortant Classification

We classified reassortants using K-means clustering according to their dispersal scales and host range. Specifically, we fitted to i) the number of different bird orders from which the reassortant had been sampled (bird richness), ii) whether the reassortant had been isolated in mammals or not, iii) the maximum geographic distance between samples within each reassortant, iv) the number of genomes, v) weighted diffusion coefficient, vi) evolutionary rate, vii) persistence time (time between the latest sampling date and the estimated date for the MRCA), vii) the number of ancestral host transitions and ix) the number of ancestral host transitions to mammals. Prior to model fitting, we normalised numeric variables and encoded presence in mammals using one-hot encoding. We iteratively fitted clusters for *K* in 1, 2, …, 20, selecting the optimal value of K according to the ‘Elbow’ method. For the optimal value of K, we inferred the relative importance of each variable to cluster assignment with a permutation analysis. Specifically, we sequentially excluded one variable from the dataset, refitted the model, and quantified the difference between each permutation and the original dataset using the adjusted rand index (ARI) [66, 67]. We implemented these models using the “stats” v4.4.2 package in R v4.1.2 [68]. Extensive use was made of the Tidyverse suite v2.0.0 for data handling, Recipes v1.09, and Broom v0.2.9.6 in the model pipeline [69, 70].

In a second statistical model, we estimated the probability that a novel reassortant, *y*_*i*_, is assigned a class, *c*, from the ordered set *C* = {minor, moderate, major}. We assumed that the probability a reassortant is assigned a given class follows a cumulative distribution, with classes increasing from minor to moderate to major [71]. We modelled each class as the discretisation of a latent (unobserved) continuous variable, 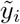, via threshold parameters, *τ*_*c*_, which partition the distribution into class-specific intervals. For each threshold, we modelled four linear predictors. Specifically, we included i) the class identity of the reassortant immediately ancestral to reassortant *i* (with respect to HA), ii) origin continent, iii) the number of segments changed relative to the immediately ancestral reassortant, and iv) the time interval between the MRCA of reassortant *i* and that for the most recent major reassortant. We modelled a penalised thin plate regression spline to smooth the time interval between the MRCA of reassortant *i* and that for the most recent major reassortant [72]. A full description of the statistical model is provided in the supplementary methods (Appendix A.2).

#### 2.5.3 Reassortant Diffusion Coefficient

To evaluate determinants of novel reassortants dispersal, we fitted a mixed model to predict the weighted diffusion coefficients (km^2^ year^-1^) [73], *y*, calculated from our phylogeographic analysis for each novel reassortant, *i*. We restricted our analysis to reassortants with a clade size greater than 1, since we cannot confidently distinguish between reassortants that truly exist at a single locus and reassortants with limited (but non-zero) circulation and incomplete sampling. For all *y*_*i*_ > 0, we assumed a gamma distribution parametrised such that, *y*_*i*_ ∼ Gamma(*κ*_*i*_, *θ*_*i*_), with rate parameter, 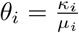, shape parameter *κ*_*i*_ and mean *µ*_*i*_.

We modelled a linear combination of predictors for each of *µ*_*i*_ and *κ*_*i*_. For *µ*_*i*_, we included predictors for the number of host-state transitions across the phylogeny, the proportion of evolutionary time (sum of branch lengths) in wild Anseriformes spp (including ducks, geese and swans), wild Charadriiformes spp (including waders, gulls and auks) and domestic Galliformes spp (including turkeys, chickens and quail); persistence time, and the number of sequences associated with each reassortant. Because the proportions of evolutionary time spent in each host group are compositional, we represented these values using isometric log-ratio coordinates [74]. We log-transformed the number of host-state transitions and the number of sequences prior to model fitting. For both *µ*_*i*_ and *κ*_*i*_, we included continent-specific random intercepts. For *µ*_*i*_ only, we allowed the effects of host composition to vary by continent, modelling continent-specific intercepts and slopes for two host-composition coordinates as uncorrelated random effects (i.e. continent is the grouping factor for non-centred random effects). Finally, we included random intercepts for the date of the reassortant MRCA, grouped by calendar year. A full description of the statistical model is provided in the supplementary methods (Appendix A.3).

#### 2.5.4 Computation

For the statistical models, we fitted all except ‘Reassortant Clustering’ to median posterior estimates obtained from haemagglutinin only, because it is the most consistently sampled and best-resolved genomic segment across the dataset, providing the most reliable estimates for further analysis (Figures S5). For the initial clustering, we fitted the model to median posterior estimates from the HA and PB2 segments. These capture complementary evolutionary signals: HA reflects antigenic and epidemiological structure, whereas PB2 captures internal-gene evolution and has well-characterised signatures of host adaptation. By using both segments, we improve the robustness of cluster assignment while avoiding biases introduced by sparsely sampled segments and introducing collinearity that can arise when analysing all eight segments simultaneously.

Due to limited prior knowledge of the relationship between the explanatory and predictor variables, we specified weakly informative priors for all Bayesian models. Exact prior specifications for each model are described in the supplementary methods (Appendix A). To fit each model, we ran four parallel MCMC simulations, each of 5000 iterations. The first 500 iterations of each chain were discarded. We assessed the convergence of posterior chains and adequate mixing against criteria of *ESS* > 1, 000 and rank-normalised Potential Scale Reduction Factor (PSRF) of 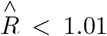 [75]. We conducted further visual checks for within-chain autocorrelation, parameter identifiability and the fit of the posterior predictive distribution to our data (Figures S8-S10, S16-S18, & S23-S25). In the main text, we report posterior predictions based on the expected value of the posterior predictive distribution. Full posterior predictive distributions, which incorporate both parameter uncertainty and stochastic variability, are included in the Supplementary Material. Throughout the manuscript, we use 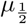 to denote the median. The ‘number of reassortants’ model was fitted in Stan using cmdstanr v0.8.1 and the remaining models were fitted in Stan using BRMS v2.20.4 in R v4.1.2 [68, 76–78]. Extensive use was made of the Tidyverse suite v2.0.0 for data handling, and tidybayes v3.0.6 and marginaleffects v0.25.0 for post-processing [70, 79, 80].

## 3 Results

### 3.1 Patterns of Novel Reassortant Emergence

We identified 209 unique reassortants across the clade 2.3.4.4b H5Nx genomes assembled for this study (Figures 1A, 1B & 1C). Importantly, these data encompass critical changes in HPAI H5 epidemiology, including the near-total replacement of H5N8 by H5N1 in 2021, the H5N1 panzootic and the initial phase of H5N5 resurgence in early 2024 (Figures S1 & S2A). Concomitantly, H5 HPAIVs expanded to a broader range of new hosts in addition to the increasing number of infections in major reservoir species (Figure S2B). This resulted in sequential epizootics in European seabirds, marine mammals in South America, and more recently, dairy cattle in North America. Reconstructing the ancestral patterns of reassortant profiles for each continent, we inferred the number of unique reassortants emerging from each (Figure 1E). Marginal posterior summaries of the number of unique reassortants originating per continent indicate Asia harboured the largest number (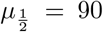 [*IQR* : 89 − 91]), followed by Europe (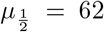 [60 63]), North America (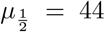 [43 − 45]) and Africa (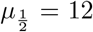 [11 − 12]). We did not infer the origins of any reassortant to be located in South America, Southern Ocean and Antarctica, or Australasia and Oceania during this time period, highlighting these regions as sinks for newly emerged reassortants that diffuse southwards from North and central America and Asia, respectively.

**Fig. 1.**
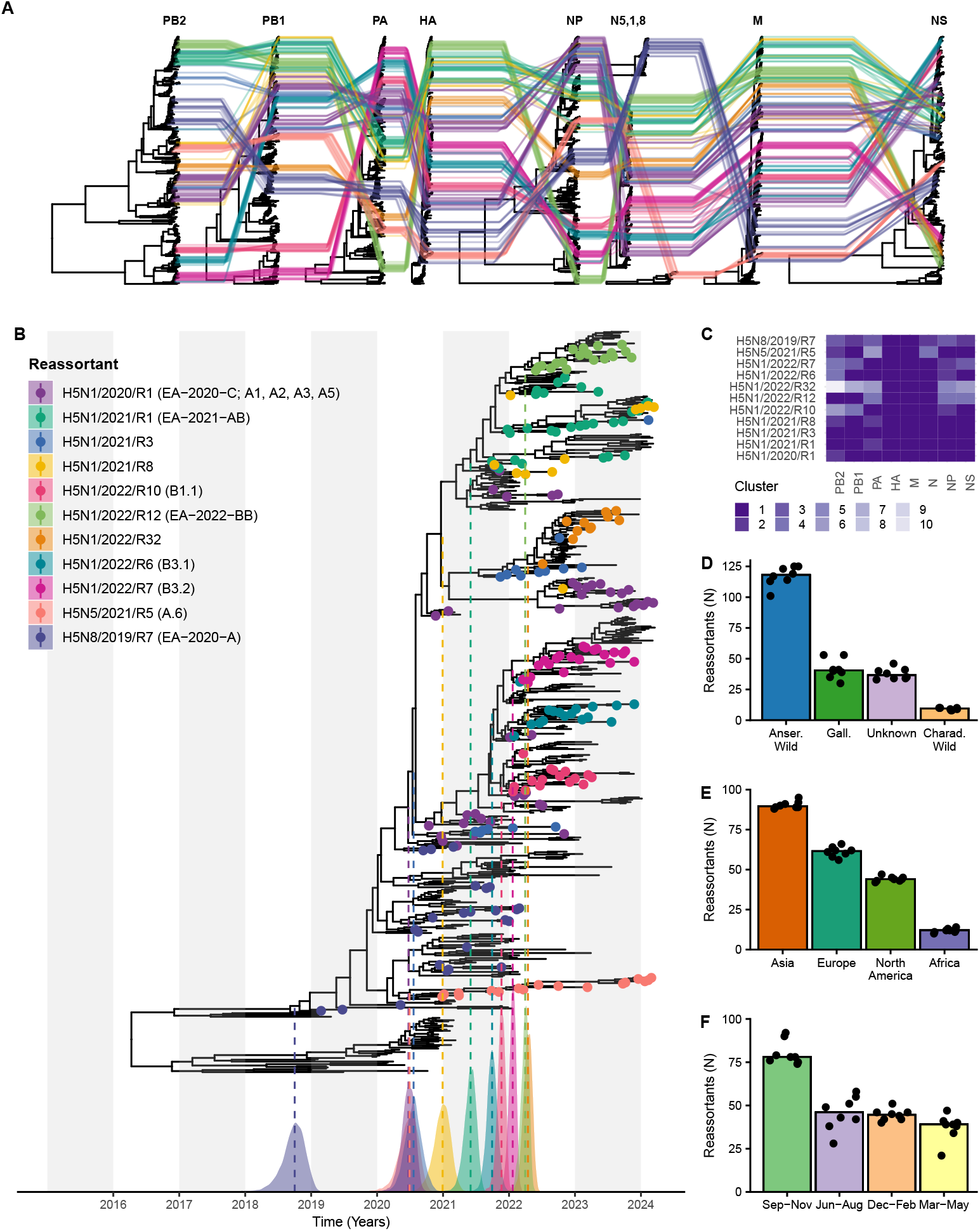
Global HPAI H5 clade 2.3.4.4b reassortment diversity between 2019 and May 2024. (A) Incongruent time-resolved phylogenies between gene-segments suggests substantial reassortment during the epizootic period 2019-2024. The coloured lines reflect linkage patterns across segments for the eleven most numerous reassortants in our data. (B) For the same eleven reassortants, we mapped estimated the time of their most recent common ancestor relative to the time-resolved haemagglutinin phylogeny. Here, coloured tips denote the unique reassortants in our dataset, assigned using patristic-distance based clusters presented in (C). The coloured densities correspond to the highest posterior density of the date of the most recent common ancestor of each reassortant, where there dotted line represents the median. Where available, each reassortant is referenced against its counter-part H5N1 genotype from the EU Reference Laboratory (EURL) and the United States Department of Agriculture (USDA). Bar plots show the number of reassortants stratified by host origin (D): wild Anseriformes (Anser. wild), Galliformes (Gall.), wild Charadriiformes (Charad. wild), and unknown (environmental or equivocal); origin continent (E); and season (F): September-November (Sep-Nov), December-February (Dec-Feb), March-May (Mar-May), June-August (Jun-Aug). Black dots show the median posterior count estimate for each segment, and the coloured bar height shows the median posterior estimate aggregated across segments.

We also compared reassortant emergence across different host groups (Figure 1D). Marginal posterior summaries indicate most reassortants emerged in wild Anseriformes spp (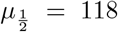 [114 124]), Galliformes spp (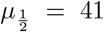 [38 − 44]), Charadriiformes spp (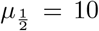 [9 10]) and other wild birds (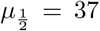 [34 − 41]). Stratifying these estimates across continents, we found wild Anseriformes spp to be the primary origin host for clade 2.3.4.4b H5Nx reassortants in Asia, Europe, and North America. In contrast, reassortants emerging from Africa were likely to have originated in domestic Galliformes; however, relative undersampling sampling in this region likely constrains the accuracy of the host origin estimation. While novel H5Nx reassortants occur year-round, these data also reveal clear temporal trends in reassortment (Figure 1F). Specifically, marginal posterior summaries indicate novel H5Nx reassortants were more often detected during September–November (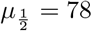 [76−82]), followed by June–August (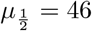 [41−52]), December–February (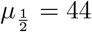 [24 − 46]), March–May (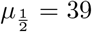 [37 − 40]).

### 3.2 Reassortant Emergence and Diversity Through Time

We quantified the rate of reassortant emergence using a hierarchical model designed to disentangle ecological and anthropogenic effects. After accounting for variation in sampling and virus ecology, we estimated a true (latent) number of reassortants per month. Assuming a fixed number of HPAIV cases and marginalising over calendar year, we found that reassortants most frequently emerged from Asia at a rate of 2.62 (95% Highest Posterior Density (HPD): 1.61 - 3.83) per month, followed by Central and North America (2.40 [1.27 - 3.77]), Europe (1.49 [0.861 - 2.28]), and Africa (0.347 [0.0972 - 0.736]) (Figure 2A).

**Fig. 2.**
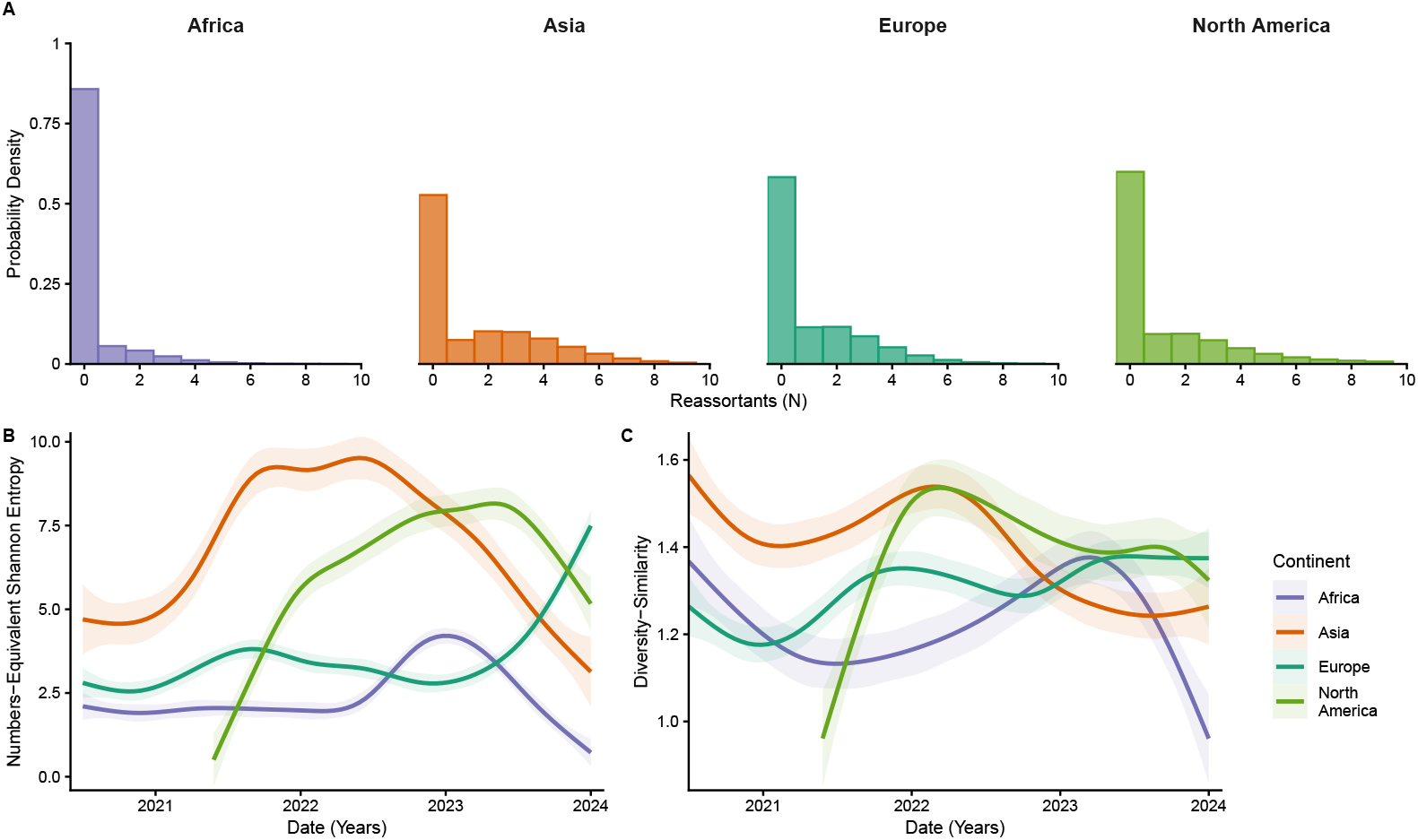
Number and diversity of H5 reassortants over time. (A) Posterior distribution of the inferred true (latent) number of reassortants, stratified by continent. (B) Numbers-equivalent diversity over time, stratified by continent. Calculated as the exponent of Shannon entropy, the numbers-equivalent diversity represents the effective number of equally likely states required to generate the estimated Shannon entropy. (C) Diversity-similarity index over time, stratified by continent. The Diversity-similarity index adjusts the numbers-equivalent diversity by genetic distance between reassortants. In (B) & (C), shaded regions correspond to 95% confidence intervals

We anticipated *a priori* that sampling effort and and prevailing environmental conditions may alter the observed reassortment frequency. To address this issue, we explicitly assumed that only a proportion of observation windows could be permissive for reassortment and that sampling of reassortments is incomplete. Our results indicate that the probability of conditions conducive to reassortment was lowest in Africa (0.564 [95% HPD: 0.151 - 0.931]), followed by Asia (0.821 [0.516 - 0.977]), Europe ( 0.824 [0.468 - 0.983]), and North America (0.905 [0.662 - 0.994]). Conditional on reassortment having occurred, an increasing number of sequences per month was intuitively associated with a higher probability that the extant reassortant is sampled. Averaged across continents, as the number of sequences per month increased threefold from 2 to 6, the probability of sampling rose from 0.504 (95% HPD:0.295-0.784) to 0.993 ( 0.941 - 1.00 ); however, the relationship between sequencing volume and sampling probability approached an asymptote, with successive increases in sequencing effort yielding a progressively smaller marginal gain(Figure S7).

Since the rate of reassortant emergence varied through time and across continents, we also sought to understand the role of viral genetic diversity in patterns of reassortment. First, we estimated the effective number of distinct reassortant profiles occurring in each continent over time using the exponent of Shannon entropy (Figure 2B). Under this approach, a diversity score of 7 implies a system with diversity equivalent to 7 distinct reassortant profiles of equal frequency. In Asia, we observed consistently high reassortant diversity, largely due to co-circulation of several reassortant types with balanced frequencies. In contrast, Europe exhibited lower diversity before 2023, with a single reassortant (H5N1/2021/R1) dominating. By late 2023 to early 2024, following the disappearance of the H5N1/2022/R12 reassortant, reassortant diversity in Europe increased, even though fewer sequences were detected. The increase in reassortant diversity can be attributed to a more balanced distribution of profiles during this period, with an absence of a single dominant type. These patterns were robust to repeatedly downsampling of regions at a uniform frequency (Figure S11).

Second, we estimated the reassortant diversity score over time based on genetic distances between reassortant profiles in each continent (Figure 2C). A higher diversity score suggests (i) more distinct reassortant profiles, (ii) more balanced frequencies, and (iii) greater genetic distances between reassortants, reflecting not only the number of reassortants but also their genetic distinctiveness. In Europe and Asia, despite temporal fluctuations in reassortant diversity, the genetic distance between circulating strains remained relatively small. This is because each new reassortant that emerged differed from the major reassortant of the previous season by only a single gene segment. Consequently, despite the observed rise in profile diversity, the overall genetic distinctiveness among these reassortants remained relatively low (Figure 2C). In contrast, North America exhibited a sharp rise in reassortant diversity between mid-2021 and mid-2022, which correlates to the introduction of European H5Nx. Subsequent reassortment occurred between the introduced European strains and the genetically distinct low pathogenicity avian influenza virus (LPAIV) gene pools. Thereafter, the reassortant diversity score decreased as local North American strains gradually outcompeted the European-originated gene segments.

### 3.3 Episodic Nature of Reassortant Emergence

We further explored the reassortment dynamics of clade 2.3.4.4b H5 HPAI by inferring the ancestral reassortant network relative to the HA time-scaled phylogenetic tree (Figure 3A). The resulting graph is highly structured and reveals an episodic pattern of reassortant emergence. Five pivotal reassortants (H5N1/2021/R1, H5N1/2022/R32, H5N1/2020/R1, H5N8/2019/R3, and H5N8/2019/R7) were the source of over half of all reassortants, each giving rise to a median of 17 (IQR: 16–28) new reassortants. Collectively, these highly connected reassortants formed a backbone that sustained the 2020–2024 panzootic across multiple seasons, facilitating global dispersal.

**Fig. 3.**
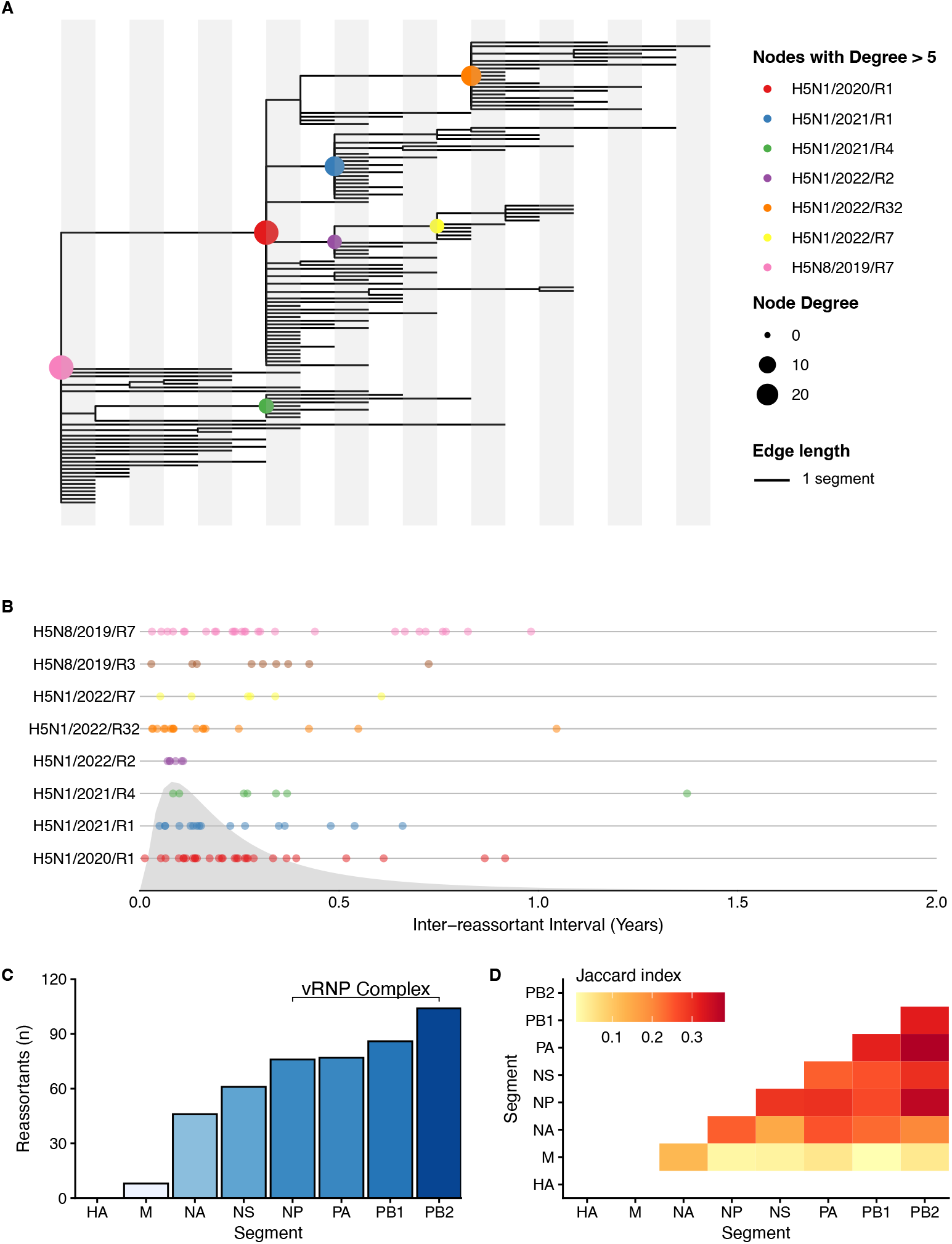
Episodic nature of H5 reassortant emergence. A) A directed network of 2.3.4.4b reassortants that emerged between 2019 and 2024, with respect to the HA phylogeny. Edge lengths are scaled by the number of segments changed between connected nodes, where each white/grey bar represents one segment. Node size is scaled by out-degree (the number of immediate offspring nodes) and nodes with an out-degree greater than five (i.e reassortants with more than five immediate descendant reassortants) are coloured (*n* = 8). B) For the eight reassortants with the most immediate descendants, we inferred the emergence time of their ‘offspring’ relative to the MRCA of the ‘parent’. Collectively, these inter-reassortant generation times followed a log-normal distribution with real-space mean 0.196 and standard deviation 0.923 (Log-likelihood = *−* 105.20), reflecting a bias towards early reassortment. Across our data, segments forming part of the viral ribonucleoprotein complex (vRNP) were exchanged with greater propensity than other segments (C). We quantified the tendency of each segment-pair (A and B) to be exchanged together using a Jaccard index 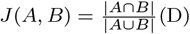. The colour and size of the dots plotted from the resulting symmetrical matrix are scaled by the magnitude of the index score

Among all reassortants with at least one ‘offspring’, the number of descendants was positively correlated with persistence time (Spearman’s Rank Correlation (SPC), *ρ* = 0.742, *p* ≤ 0.001), intuitively suggesting that longer circulation increases the likelihood of further reassortment. Under this scenario, one might also expect the inter-reassortant generation time to increase concomitantly with persistence time; however, in our dataset no correlation was present (SPC, *ρ* = 0.00, *p* = 0.9861) with a median inter-reassortant generation time of 3 months and 26 days (Figure 3B). Most reassortants arose early in the lifespan of their ‘parent’ reassortants, possibly due to initial ecological opportunities or the higher fitness of early variants, rather than accumulating through prolonged circulation.

To investigate patterns of individual gene exchange, we mapped changes in genome composition from one reassortant to the next across the network. Segments comprising the viral ribonucleoprotein (vRNP) complex were exchanged most frequently (Figure 3C). Half of all reassortants (*n* = 104) acquired a new PB2 segment relative to their immediate ancestor, followed by PB1 (*n* = 86), PA (*n* = 77), and NP (*n* = 76), which were the second, third, and fourth most frequently exchanged segments, respectively. An exact binomial test showed that the number of reassortants where all vRNP segments were replaced (17 out of 209) was significantly greater than expected by chance (*p <* 0.001, one-tailed). To identify segment pairs that frequently reassorted together, we calculated the ratio of the intersection to the union (Jaccard Index) of segment-switching events. This analysis revealed a propensity for internal segments to reassort in combination (Figure 3D). Notably, PA and PB2 were significantly more likely to be exchanged together than individually (permutation test, *p* = 0.042). We also identified a significant, though less frequent, association between the reassortment of NA and M segments (*p* = 0.028).

Finally, we recorded differences in the evolutionary contribution of reassortments between host class. Specifically, reassortants for which a median of 80% of isolates across countries were detected in domestic Galliformes (*n* = 36) were significantly less likely to be ‘parent’ reassortants than those with a median ≥ 80% of isolates in wild Anseriformes (*Z* = 2.21425, *p* = 0.0134) (Figure S12).

### 3.4 Phylodynamic Reassortant Classes

To characterise observed differences between reassortants, we clustered each of the 209 reassortant according to their phylodynamic profiles. These data included posterior estimates of evolutionary rate, host-state transition rates, diffusion coefficient, and persistence times. We identified three main clusters of reassortant, corresponding to levels of circulation across spatio-temporal, ecological, and epidemiological scales (Figure S14). Specifically, group A (minor) comprised 180 (86.12 %) tightly clustered reassortants profiles (median pairwise Euclidean distance = 1.98), group B (moderate) comprised 24 (11.48%) reassortants, and group C (major) comprised five (2.39%) weakly clustered reassortants (median pairwise Euclidean distance = 11.44). We calculated that ‘minor’ reassortants generally persisted for a median of 0.209 years (IQR: 0.0897 – 0.499), rarely switched hosts (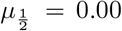; IQR: 0.00 – 0.00), and included at least one mammalian sample in negligible proportion of cases (0.060; 95% CI: 0.031 – 0.110). In contrast, ‘moderate’ reassortants circulated for a median 1.24 years (IQR: 0.426 - 1.94), which coincided with semi-regular host-state transitions (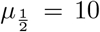; IQR: 0.00 - 19). Approximately half (0.541, 95% CI: 0.332 - 0.738) of ‘moderate’ reassortants included samples obtained from mammals. Finally, ‘major’ reassortants circulated the longest, with a median persistence time of 3.08 years (IQR: 2.25 - 3.78). All ‘major’ reassortants included samples obtained from mammals (95% CI: 0.462 - 1.00) and with a median of 215 host switches each (IQR: 145 - 292) (Summarised in Figure 4A).

**Fig. 4.**
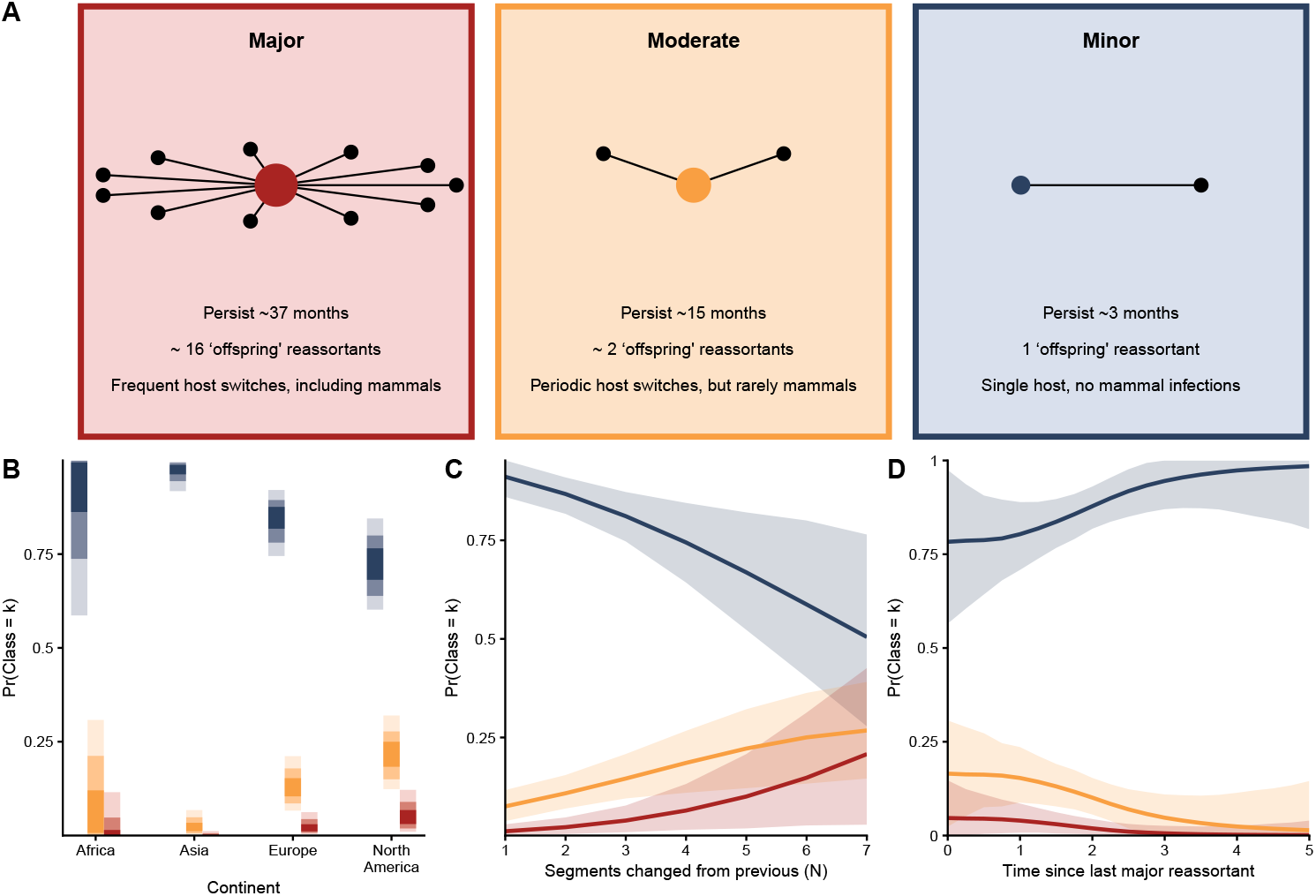
Reassortant Class. (A) Summary of reassortant classes with defining characteristics. (B) Posterior estimates of the probability that a given reassortant is of class, k, stratified by continent. Opacity indicates highest posterior densities for each estimate, with increasing opacity corresponding to the 95%, 85%, and 50% thresholds. The average marginal effect of the number of segments exchanged (C) and the time since the last major reassortant (D) on the reassortant class probability. (D) The ‘last’ reassortant is defined according to the ancestral state reconstruction (ie a phylogenetic order rather than chronological order). For both C and D, the shaded area indicates the 95% highest posterior density.

Out of the five major reassortants, we estimated that three emerged in Europe (H5N1/2020/R1, H5N1/2021/R1, and H5N1/2022/R12), one in Africa (H5N8/2019/R7), and one in North America (H5N1/2022/R7). Consistent with previous studies, our phylogenetic analysis across multiple segments showed H5N1/2020/R1 was formed from the reassortment of H5N8/2019/R7 segments HA and MP with LPAIV circulating in Europe [20]. We identified H5N1/2020/R1 as an ancestral variant of almost all subsequent clade 2.3.4.4b reassortants during 2020-2024 (Figure S15). The following season, our phylogenetic analysis showed H5N1/2020/R1 exchanged segments PB2 and PA with LPAIV H5N3 to form H5N1/2021/R1, which we also classified as a dominant reassortant. Throughout 2022-24 our phylogenetic analysis reconfirmed H5N1/2021/R1 had acquired new PA, NP, NS segments from LPAIV H13, giving rise to H5N1/2022/R12. Unusually, this major reassortant circulated extensively during the summer of 2022, with a specific burden on Charadriiformes spp [81, 82]. We found H5N1/2022/R12 was the ‘major’ reassortant with the lowest number of ‘offspring’ reassortants (*n* = 3), all of which were minor with very limited circulation.

Our phylogeographic analysis reconfirmed H5N1/2020/R1 within wild birds facilitated dispersal from Europe to North America through migratory routes over the Atlantic Ocean [83]. Colonisation of North America exposed Eurasian clade 2.3.4.4b HPAIVs to locally circulating LPAIVs, triggering a new wave of reassortment between the two. In this study, we identify major reassortant H5N1/2020/R1 was ancestral to the first ‘North American’ reassortant, H5N1/2022/R2, which subsequently exchanged PB2 and NP segments before undergoing further reassortment exchanging PB2, PB1 and NS. We classified the resulting reassortant, H5N1/2022/R7, as major. Consistent with previous studies, our phylogeography revealed H5N1/2022/R7 dispersed widely across northern and South America causing repeated spillovers into terrestrial and marine mammals, finally reaching the Antarctic region by October 2023.

Moderate reassortants also played a pivotal role in shaping the evolution of clade 2.3.4.4b H5, albeit at a smaller scale relative to major reassortants. For example in our data, reassortant H5N1/2021/R3 comprised 70% of genomes isolated from wild birds in China since 2021, having originated from wild birds in southern Africa in the autumn of 2020. Our phylogeographic analysis showed H5N1/2021/R3 spreading among wild migratory birds in China and also infected poultry and wild birds in eastern (mainly South Korea and Japan) and south-eastern Asia during the 2022/2023 season, even leading to human infections in eastern China in 2023. Similarly, H5N1/2023/R29 (B3.13) is the moderate reassortant responsible for the initial outbreak in US dairy cattle and over 20 subsequent human infections [84–86].

Notably, we identified six moderate reassortants had five or more ‘offspring’ reassortants (H5N1/2022/R32 (17), H5N8/2019/R3 (9), H5N1/2021/R4 (7), H5N1/2022/R2 (6), H5N1/2021/R3 (5) and H5N1/2023/R6 (5)), highlighting the capacity of these ‘moderately fit’ variants to further influence viral evolutionary dynamics.

Next, we analysed whether patterns of the frequency of specific reassortant clusters varied through time and by continent. Marginalised over the empirical data distribution, we estimated that ‘major’ reassortants were proportionally most likely to have originated in North America (with probability 0.057 [95% HPD: [0.011 - 0.121]), followed by Europe (0.026 [0.003 - 0.062]), Africa (0.014 [0.003 - 0.062]), and Asia (0.003 [0.000 - 0.012]) (Figure 4B). Similarly, we found that moderate reassortants were also more likely to originate in North America (0.215 [0.120-0.314]) and Europe (0.134 [0.067 - 0.210]) than either Africa or Asia (0.014 [0.00 - 0.099] and 0.003 [0.00 - 0.012], respectively). Perhaps unsurprisingly, we found minor reassortants occurred at high frequency across all continents. For all reassortant classes, our predictions for Africa were least certain, likely due to reduced sampling relative to other continents. Our posterior estimates were robust to sensitivity analysis exploring a counterfactual scenario where the proportion of wild bird sequences was fixed (at 0.25, 0.5, and 0.75) for each region (Figure S19).

Averaged across the empirical data distribution, major and moderate reassortants were most likely to follow a major reassortant with probabilities 0.035 (95% HPD: 0.008 - 0.071), and 0.143 (0.084 - 0.209), respectively. Specifically, H5N8/2019/R7 is immediately ancestral to H5N1/2020/R1, and H5N1/2020/R1 is itself immediately ancestral to H5N1/2021/R1 and H5N1/2022/R12. H5N1/2022/R7 is the only major reassortant not immediately descended from another major reassortant, instead inter-spersed by moderate reassortant H5N1/2022/R2. If a minor reassortant was ancestral to any reassortant at all, it was highly likely to also be minor (0.946 [0.873 - 0.995]).

We also identified differences in the reassortant class-identity according to the number of new segments relative to the ‘parent’ reassortant and the time since the last major reassortant. If two segments were exchanged relative to the parent reassortant, we estimated the probability of a major reassortant to be 0.0223 (95% HPD: 0.004 - 0.047), which increases three-fold when the number of segments exchanged was raised to four (0.063 [0.016 - 0.128]) (Figure 4C). Marginalised over the empirical data distribution, the average affect of each additional segment increased the probability of a reassortant being ‘moderate’ or ‘major’ by 3.84 (1.21 - 6.84) and 1.67 (0.251 - 3.79) percentage points, respectively. Finally, at shorter inter-major-reassortant intervals, the probability of major and moderate reassortants was relatively high (0.0446 [0.006 - 0.109] and 0.166 [0.076 - 0.272], respectively), decreasing rapidly as the interval increased (Figure 4D). Conversely, the probability of a minor reassortant increased with time increasing from 0.786 (0.650 - 0.9145) at a six month interval to 0.944 (0.869-0.998) with a 3-year interval.

### 3.5 Estimating The Reassortant Diffusion Velocity

To understand factors associated with the circulation of emerging reassortants, we inferred the dispersal history of all reassortant profiles for which we obtained two or more spatially distinct whole genomes (Figure S6). Next, we fitted a generalized linear mixed model to disentangle the effects of host richness, relative proportions of evolutionary time in wild Anseriformes spp, wild Charadriiformes and domestic Galliformes spp and persistence time on the weighted diffusion coefficients (an estimation of the area invaded by viral lineages per unit of time) for each reassortant (Figure 5).

Marginalising over all observations in our data, the median predicted diffusion coefficient was 1,864 km^2^ day^-1^ (95% Highest Posterior Density (HPD): 916 − 3, 439 km^2^ day^-1^). We identified substantial variation in both the baseline diffusion coefficient and its heteroskedasticity across the 4 regions included in the model 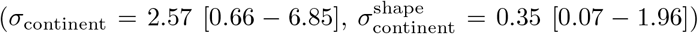. We estimated that reassortants emerging from North America diffused fastest (2,588 km^2^ day^-1^ [1, 197 − 4, 899 km^2^ day^-1^]), followed by reassortants emerging from Africa (2,327 km^2^ day^-1^ [448 − 9, 807]) and Asia (2,224 km^2^ day^-1^ [1, 029 − 4, 528]). Reassortants that emerged from Europe were predicted to be the slowest to occupy new geographies (1,506 km^2^ day^-1^ [585 − 3, 569]). We also identified moderate temporal variation in the diffusion coefficient, relative to the global baseline (*σ*_year_ =0.81 [0.10 − 2.18], Figure S20). Specifically, reassortants that emerged in 2019 had the highest average marginal diffusion coefficient (3,581 km^2^ day^-1^ [396 − 16, 274]), decreasing as the H5 HPAIV zoonotic progressed (1,315 km^2^ day^-1^ [471 − 2, 848] in 2023).

Next, we investigated the effect of host-state on the magnitude of the weighted diffusion coefficient. We estimated that the highest number of host-state transitions amongst all species per 100 whole genomes was in Southern America (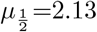 [IQR:1.70 − 2.13]), followed by Central and North America (2.11 [1.96 − 2.21]), Africa and Europe (0.683 [0.505 - 0.761] and 0.256 [0.229 − 0.279], respectively) (Figure S22). Applying these data to predict the diffusion coefficient, our fitted hierarchical model revealed that greater cross-species diversity is strongly positively associated with weighted diffusion coefficient (1.75, 95% HPD: 0.67–2.85). By averaging across each combination of unique categorical predictors while holding numeric variables at their means; we estimated that all else equal the weighted diffusion coefficient increases from 1,595.36 km^2^ day^-1^ (95% HPD: 803.53 − 2, 850.12) for 1 host-state transition, to 12,402.22 km^2^ day^-1^ (1, 370.42 − 50, 186.46) with ten host-state transitions (Figure 5B). The number of sequences per reassortant also positively correlated with the weighted diffusion coefficient (0.87 [0.11 − 1.65]), but it did not concomitantly increase the number of host-states.

**Fig. 5.**
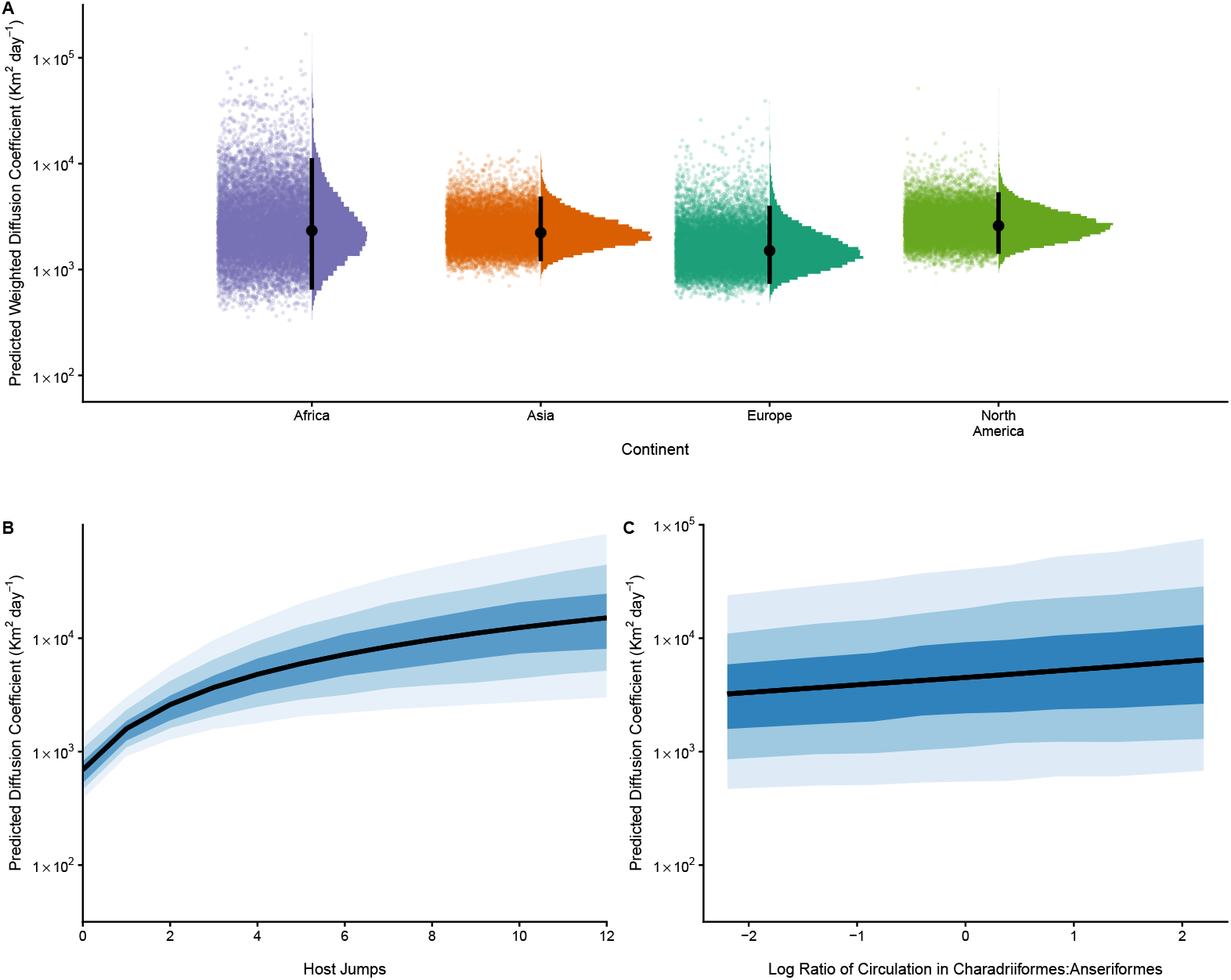
Diffusion coefficient estimates of H5Nx reassortants stratified by time, location, and host persistence. (A) Expected value of the posterior predictive distribution for the weighted diffusion coefficient, stratified by continent of origin. Each draw represents the mean prediction conditional on a given set of posterior parameter values, summarising the central tendency of the model predictions. (B) Posterior predictive distributions of the diffusion coefficient as a function of the number of host-state transitions and (C) the proportional circulation of Charadriiformes vs Anseriformes. Each is marginalised over each combination of unique categorical predictors and the mean of all numeric variables. Shaded regions indicate highest posterior density intervals for each estimate, corresponding to the 95%, 85%, and 50% levels from light to dark

Our model also revealed subtle variation between the diffusion coefficient estimated for each reassortant and the relative persistence in select bird orders. Specifically, we estimated that reassortants with a proportionally greater persistence in Charadri-iformes than Anseriformes show weakly enhanced diffusion (0.43, HPD: 0.01-1.58) (Figure 5C). For a fixed persistence of one year and averaged over all other variables, the weighted diffusion coefficient for a reassortant with a 1:1 ratio of circulation in Charadriiformes to Anseriformes was 4,505.21 km^2^ day^-1^ (146.89 − 31, 075.91). As the relative proportion of Charadriiformes to Anseriformes increases, the predicted weighted diffusion coefficient also increases. For example, when viral circulation in Charadriiformes is twice that in Anseriformes, the expected diffusion coefficient of an average reassortant persisting for one year increases to 5,157.39 km^2^ day^-1^ (197.43 − 38, 510.85). By allowing the effect of the relative proportion of Charadriiformes to Anseriformes to vary across continents, we also revealed moderate region-level variation (*σ*_ilr3_ = 0.50 [0.07 − 1.96]). In our model, we also evaluated the effect of circulation in Anseriformes and Charadriifomes relative to circulation in Galliformes, and between Anseriformes, Charadriifomes and Galliformes, relative to all other hosts. In either case, the fitted coefficients did not significantly differ from zero (Figure S21).

### 3.6 Global Drivers of Reassortant Spread

Next, we investigated the role of eight environmental drivers on the spread of the five major H5Nx reassortants between regions (Figures 6 and S27). Using the GLM extension of discrete trait phylogeography, the number of shared flyways was identified as a strongly supported positive predictor of interregional virus spread in the combined five major reassortants (*β* 1.83 [0.76, 3.13]; Bayes factor (BF) > 20), as well as for the individual reassortants H5N8/2019/R7 (1.86 [0.88, 3.17]; BF > 20), and H5N1/2020/R1 (1.10 [0.54, 1.67]; BF > 20), both of which exhibit more widespread spatial patterns globally compared to the other three major reassortants. We also found that geographic distance between regions was negatively correlated with dispersal for the combined dataset (-1.04 [-1.25, -0.82]; BF > 20) and for two major reassortants H5N1/2020/R1 (-0.93 [-1.13, -0.73]; BF > 20) and H5N1/2022/R7 (-1.26 [-1.63, -0.88]; BF > 20), respectively. This would suggest diffuse and local circulation between neighbouring regions as a driving force behind the spread of these reassortants, alongside long-range migratory movements.

Across all five major reassortants, we identified duck density (0.60 [0.41, 0.79]; BF > 20) as a positive correlate with virus dispersal, re-emphasising the important role that the wild bird/domestic poultry interface plays in maintaining and propagating viral circulation. Similarly, we estimated mean annual temperatures at origin and destination locations as negative (reassortant H5N8/2019/R7; 0.73 [0.25, 1.44]; BF > 20) and positive (five major reassortants combined; -0.60 [-0.94, -0.33]; BF > 20) predictors of virus dispersal, respectively. This pattern may reflect that higher temperatures could increase food availability and productivity, thereby attracting migratory birds that serve as potential carriers for viral dissemination.

**Fig. 6.**
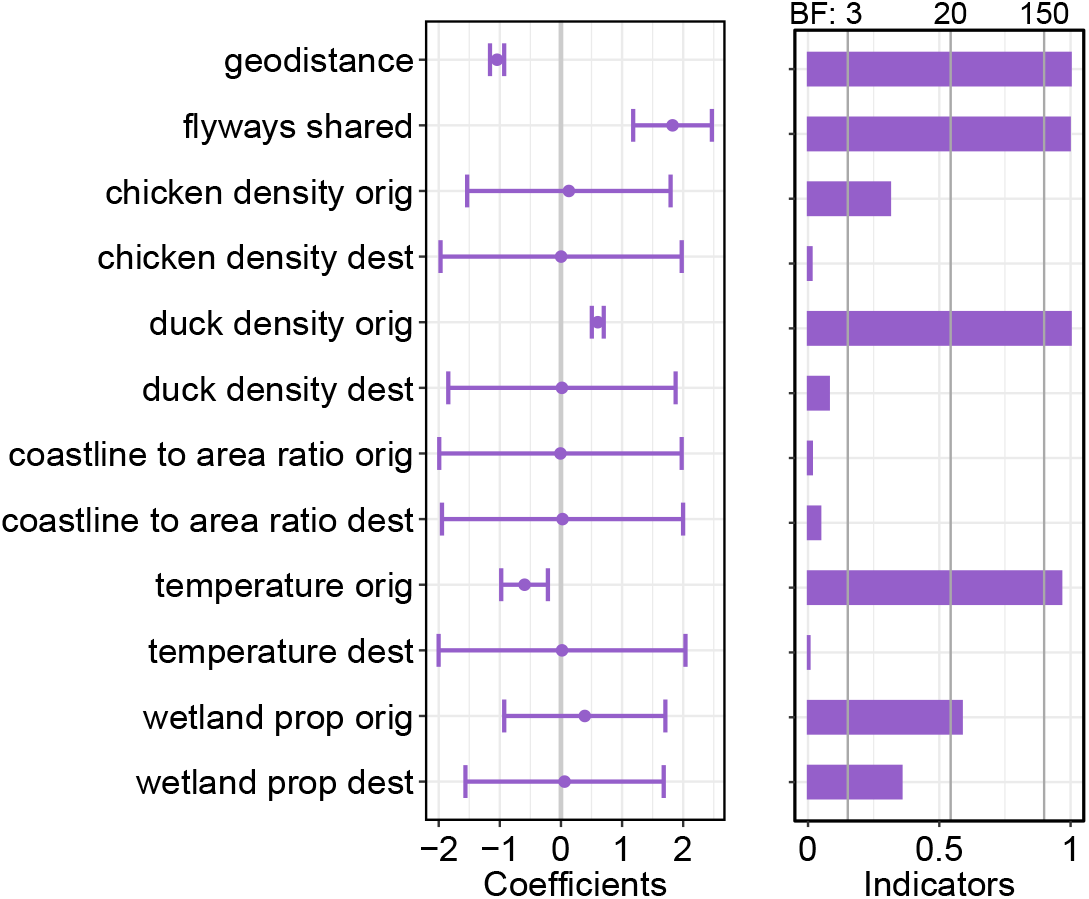
Contributions of predictors to the interregional spread of five dominant H5Nx AIV reassortants. We inferred virus dispersal patterns by extending a discrete phylogeographic inference with a phylogeographic generalised linear model (PGLM). Where selected predictor variables were highly correlated (absolute Pearson correlation coefficient > 0.66), we repeated the analysis separately for each predictor. The left panel shows the posterior means and 95% highest posterior densities of the model coefficients, which quantify the effect size of the predictor (in log space) on the region-state transition rates. The right panel shows the posterior inclusion probability of each predictor variable, and their corresponding Bayes factor (BF; grey solid vertical lines in right panel), which quantifies their statistical support for inclusion in the model. BF *<* 3: no or weak support; 3 *≤* BF *<* 20: support; and BF *≥* 20: strong support [87].

For both H5N1/2021/R1 and H5N1/2022/R12, we did not identify any statistically supported predictors of virus spread and the fitted regression coefficients did not deviate significantly from their prior distributions. The circulation of both H5N1/2021/R1 and H5N1/2022/R12 was largely restricted to Europe, where the relative homogeneity of environmental and anthropogenic landscapes could reduce the explanatory power predictors aggregated at the global scale. Further analyses conducted at finer geographic scales may reveal additional drivers underlying the spread of these reassortants.

## 4 Discussion

The resurgence of clade 2.3.4.4b coincided with step-changes in the evolutionary dynamics of H5 HPAIVs. In 2020, novel reassortant H5N1/2020/R1 not only out-competed contemporary HPAIV H5N8 lineages, but triggered the start of a global epizootic and ecological crisis in birds and marine mammals [27]. Thereafter, pervasive reassortment has generated a vast array of novel genetic combinations, concurrent with an expansion in host range, long-term persistence and worldwide dispersal [18]. In this study, we systematically characterised global patterns of reassortment, identifying 209 unique H5Nx reassortants circulating during the 2020-2024 panzootic. We revealed a structured, intermittent pattern of reassortant emergence, in which five ‘major’ reassortants dominated. We estimated that a greater number of reassortants originated from Asia, while reassortants from North America spread most quickly. Finally, we reconfirmed the critical role for wild bird movement in H5 HPAIV dispersal and persistence. Our results suggest that Charadriiformes spp may subtly increase the diffusion coefficient of newly emerged reassortants; however, we also show that diffusion appears more sensitive to opportunities for cross-species transmission than to the specific host-order composition.

Across the 2020-2024 panzootic, we characterised an episodic pattern of reassortant emergence, in which most new reassortants appeared shortly after their direct ancestors irrespective of the overall persistence of the segment ‘donors’ (Figure 3B). This transient peak in reassortant emergence is consistent with a general invasion dynamic, whereby reassortment is limited to periods of coexistence before competitive displacement reduces opportunities for coinfection. Several overlapping ecological processes may widen or narrow this coexistence window. First, spatial and temporal overlap among key host taxa may temporarily increase opportunities for co-infection and reassortment. During overwintering, breeding, or migration stopovers, otherwise segregated populations are more likely to intermix. These host communities may differ in immune history or competence, creating temporary “mixing vessel” conditions in which multiple viral lineages co-circulate at sufficient prevalence for reassortment [88–90]. Second, the duration of this coexistence window may be constrained by competitive interactions between LPAIV and HPAIV. Localised HPAIV outbreaks often coincide with decreases in LPAIV circulation [91]. If both viruses compete for the same susceptible hosts, invasion by HPAIV may rapidly reduce lineage overlap through immune cross-protection, mortality, or stochastic extinction, limiting opportunities for coinfection. The extent of this coexistence window likely depends on the relative fitness of competing viruses and the strength of immune cross-protection [92, 93]. Previous *in vivo* studies of Anatidae spp and *Branta canadensis* suggest prior infection by the same subtype is likely to prevent infection altogether, but prior infection by a different subtype (e.g H1N1 or H3N8) provides variable levels of incomplete protection, which may prolong infection and influence reassortment opportunity [94–99].

In this study, Asia was the source of the greatest number of reassortants, but all else equal, reassortants emerging from Asia were also the least likely to be ‘Major’ (Figures 2 & 4). Circulation of AIVs in Asia, and in particular mainland China, is closely linked to intermittent mixing of domestic and wild Anseriformes spp that facilitate cryptic circulation [97]. Given a broad standing viral genetic diversity of LPAIVs in this region, we anticipate interactions at the wild-domestic interface may transiently concentrate diverse viral lineages within temporary “mixing vessel” populations, increasing the probability of productive co-infection and the generation of novel reassortants [100]. Incidentally, this same standing diversity may also attenuate reassortant success by intensifying competition with well-adapted parental strains. Therefore, although reassortment events may arise more frequently in, the ‘success’ of emerging reassortants may be limited. Conversely, major reassortant H5N1/2022/R12 was concentrated in Charadriiformes spp, yet produced relatively few descendant reassortants compared to others of the same magnitude (Figure 3B). In contrast to other wild bird populations, European Charadriiformes spp were immunologically naïve to H5 prior to 2022, with gull-associated LPAIV subtypes H13 and H16 providing minimal immune-overlap with H5 [88, 93, 101]. In this context, invasion into a naïve host population may have accelerated the competitive exclusion of cocirculating LPAIV, potentially shortening the window of lineage coexistence and limiting opportunities for further reassortment.

In previous H5Nx epizootics, the migration Anseriformes spp has been instrumental in sustaining H5Nx circulation and extending its geospatial range [73, 102]. In this study, our PGLM analysis revealed a significant positive association between interregional state transitions and the number of intersecting terrestrial flyways, reconfirming these migratory pathways act as a driving force for AIV dispersal [21]. Nonetheless, while Anseriformes spp remain the canonical reservoir host of avian influenza, other bird orders may also facilitate disease spread [103]. For example, in North America between 2008 and 2018, Charadriiformes spp were associated with the accelerated migration of H5Nx, with dispersal rates exceeding either wild Anser-iformes spp or domestic Galliformes spp [93]. In this study, we identify a potential role for Charadriiformes to accelerate the dispersal of H5Nx relative to Anseriformes (Figure 5). In tandem with our PGLM analysis, our findings suggest that while Anseriformes continue to have an outsized role in the maintenance of viral persistence, Charadriiformes spp may contribute disproportionately to long-distance spread through rapid interregional dissemination.

The contribution of domestic birds to the evolution and spread of clade 2.3.4.4b reassortants between 2020-2024, however, is less clear. In preceding HPAI H5Nx lineages, dispersal was more closely associated with poultry systems, in which the production and intermixing of domestic ducks and landfowl presented frequent opportunities for reassortment [104, 105]. While our study also identified duck density as a positive correlate with the frequency of interregional movement of the five H5 reassortants (Figure 6), we did not identify any clear association between interregional movement and chicken density, nor the rate of reassortant diffusion and the relative proportion of persistence in domestic Galliformes spp (Figure S21). Our findings are consistent with fine-scaled local studies, which report a positive correlation between the density of active duck farms and between-farm transmission but not for the density of active chicken farms [106]. Likewise, we rarely observed reassortment between chicken-adapted AIV subtypes and 2020-2024 clade 2.3.4.4.b H5 HPAIVs, consistent with the 2016-2017 H5Nx HPAI epizootic [16]. Previous *in situ* measurements of reassortment between H5N1 and chicken-specific AIVs highlighted relatively low rates of reassortment, not attributable to high mortality of H5 HPAIVs [7]. Contemporary surveillance indicates that H5N1–H9N2 reassortants are rarely detected across eastern and southeastern Asia, despite the endemicity of H9N2 in intensive chicken production systems that could facilitate HPAIV spread [107–111]. These results may reflect distinct roles of different types of domestic poultry in viral evolution and dispersal. For example, clade 2.3.4.4b H5 HPAIVs are thought to show strong host preferences for wild Anseriformes and comparatively poor transmissibility in Galli-formes spp [112]. Similarly, reassortment between H5 and H9N2 may be constrained due to restricted genetic compatibility [113], producing reassortants with attenuated transmissibility [114].

Throughout this study, uncertainty in data sampling and coverage may bias our conclusions. First, regional variation in sentinel avian influenza surveillance means some reassortments in less-frequently monitored areas may be under-represented, and we anticipate that reactive sequencing efforts, stood-up across the globe in response to the escalating panzootic, likely exacerbated this pattern. In our study, severe under-sampling could mean a distant viral lineage is misclassified as a novel reassortment because evolutionary intermediates went unsampled. Similarly, analyses of host class switching are highly sensitive to both sampling intensity and specificity, which also likely varies between countries and/or regions. Our statistical modelling, however, emphasises the importance of the evenness of sampling across regions to capture global patterns of reassortant emergence.

Second, a natural consequence of our global perspective is a loss of spatial resolution, which limits our ability to resolve fine-scale transmission dynamics, potentially inflating dispersal estimates and obscuring local heterogeneity [115]. The current PGLM analyses similarly rely on time-static predictors and transmission metrics aggregated at region level, and therefore may not capture temporal variation, individual-level host movement (particularly in wild birds), or fine-scale spatial processes. The role of factors such as bird migration in H5 spread warrants further investigation using higher-resolution surveillance data and an improved that precisely captures inter-regional avian movement and contacts. Finally, we recommend a conservative interpretation of the continent-specific zero-inflation parameters in our ‘number of reassortants’ model which were only weakly identifiable from this data [116].

In summary, this study provides valuable insights into the mechanisms driving reassortment emergence and success, providing a comprehensive analysis of global patterns of reassortants within clade 2.3.4.4b H5 HPAIV between 2020–2024. Reassortment emerged as a key evolutionary feature of the panzootic: five major reassortant constellations achieved intercontinental spread and ultimately founded almost all subsequently circulating lineages. Notably, reassortants that spread most rapidly were reported across a broader host range, suggesting that successful global dissemination may be facilitated by increased opportunities for cross-species transmission and maintenance in diverse avian communities. In addition, a short inter-reassortment generation time appeared to promote temporally and phylogenetically clustered emergence of novel genotypes, consistent with repeated reassortment events occurring over short timescales during intense transmission. Therefore, alongside geographically balanced sequencing efforts to better resolve global evolutionary patterns of AIV, intensified surveillance remains essential to capture the full spectrum of reassortant diversity. Improved global collaboration, data sharing, and more even surveillance coverage will be critical to reduce blind spots in regions and host populations where transmission might otherwise go undetected, enabling cryptic spread and delayed recognition of emergent lineages. Ultimately, expanding surveillance capacity and promoting a more equitable approach to data collection will enhance our understanding of global avian influenza evolution and transmission.

## 5 Acknowledgements

This work has made use of the resources provided by the Edinburgh Compute and Data Facility (ECDF) (http://www.ecdf.ed.ac.uk/) and has been prepared using information supplied by the European Union’s Copernicus Climate Change Service (doi: 10.24381/cds.f17050d7). Neither the European Commission nor the European Centre for Medium-Range Weather Forecasts is responsible for any use that may be made of the Copernicus information or data it contains. We gratefully acknowledge all data contributors, i.e., the Authors and their Originating laboratories responsible for obtaining the specimens, and their Submitting laboratories for generating the genetic sequence and metadata and sharing via the GISAID Initiative, on which this research is based. We acknowledge BirdLife International for sharing their spatial data for the eight terrestrial flyways. We thank PhD students, Kunpeng Yuan and Lixia Wang; and postgraduate students, Haoyu Wen, Jia Dong, and Mingjia Wu, for their contribution to data curation.

## 6 Data Availability

All sequence data supporting the findings of this study were provided by the GISAID EpiFlu database (https://www.gisaid.org), with the GISAID Isolates tabulated in Table S1.

## 7 Code Availability

Beauti XMLs and R/Stan code pertaining to data extraction, reassortment clustering and statistical models fitting are publicly available under GPL-3.0 license at https://github.com/fLuLab/AIV2344bEpizooticReassortment.

## Appendix A Supplementary Methods

### A.1 Number of Reassortants

For each year-month observation, *i* ∈ {1, 2, …, *I*}, taken in continent, *j* ∈ {africa, asia, americas, europe}, let *y*_*ij*_ ∈ ℤ_≥0_ be the observed number of reassortants. We assume *y*_*ij*_ can be modelled as a mixture of three components: a sampling model, an abundance model, and a zero-inflation model.

#### A.1.1 Sampling Model

First, we consider that only a proportion, *p*_*ij*_ ∈ (0, 1), of true (latent) reassortants, 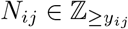, are ultimately observed:

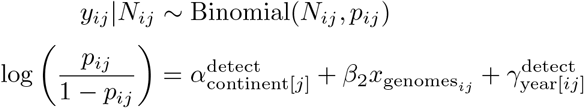

where 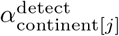, is the continent-stratified proportion of reassortants detected and 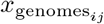, is the log-scale quantity of HPAIV full genomes present on GISAID. We assume that the interval between sequence collection and the most recent common ancestor of each reassortant is of sufficiently short duration that no lag is required to be accounted for. We specify the following weakly informative prior distributions for the sampling model:

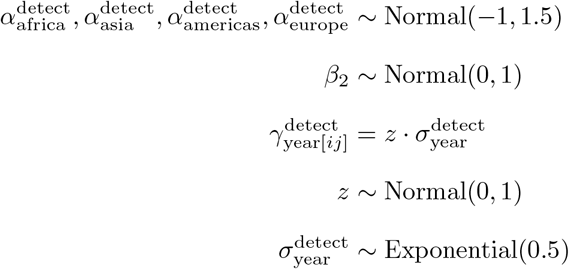

#### A.1.2 Abundance Model

Second, we model the true number of reassortants, 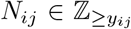, as a discrete latent variable that follows a Poisson distribution:

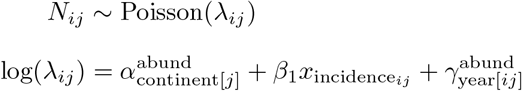

where *λ*_*ij*_ is the expected number of reassortants per observation on the log scale, 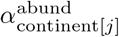, is the continent-stratified baseline abundance, 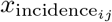, is the log-scale HPAIV incidence estimate, and 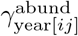 and 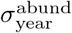 are the zero-centred random intercepts and standard deviation of calendar year. We specify the following weakly informative prior and hyperprior distributions for the abundance model:

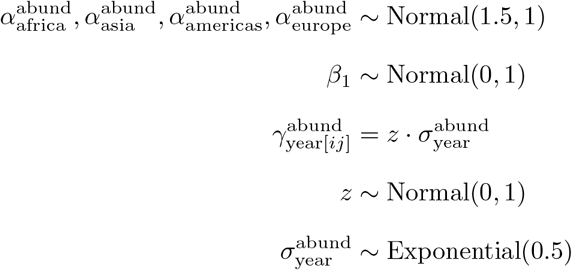

#### A.1.3 Zero-Inflation Model

Third, we consider that ecological or epidemiological conditions may not always be conducive for reassortment/reassortant emergence. We assume this process is fundamentally distinct from a structural absence of reassortment (i.e situations where reassortment/reassortant emergence is feasible but does not occur). We model a Bernoulli zero-inflation component, *z*_*ij*_ ∈ {0, 1}, parametrised by a continent-specific probability that a conditions are not permissive for reassortment/reassortant emergence, *θ*_continent[*j*]_:

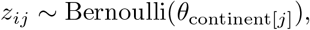

We specify the following weakly informative prior distributions for the zero-inflation model:

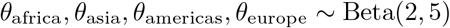

#### A.1.4 Joint Probability

We combined the abundance, sampling and zero inflation components to calculate the joint probability that *y* reassortants are recorded each month *i* per continent *j*s:

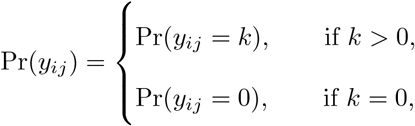

where,

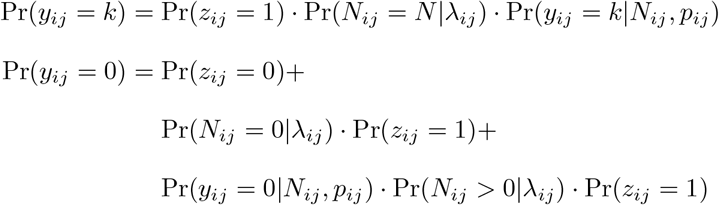

We fit this model using the probabilistic programming language Stan [78]. Since the Hamiltonian Monte Carlo sampler used by Stan cannot natively handle discrete latent variables, we integrate the joint probability over plausible values of *N*_*ij*_ ∈ {*y*_*ij*_, *y*_*ij*_ + 1, …, *y*_*ij*_ + *K* − 1} where *K* = 12 [117]:

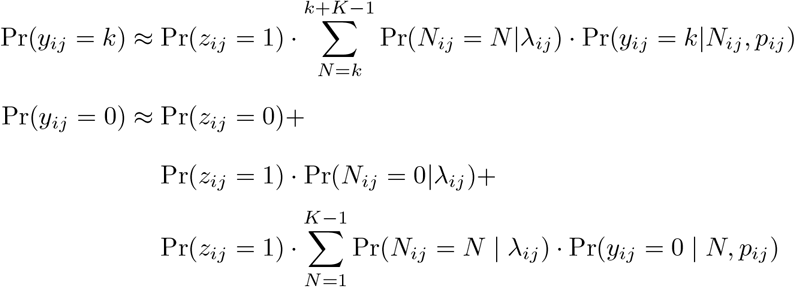

Essentially, our model describes a thinned zero-inflated Poisson distributed variable. To avoid the necessary computational approximations, one could alternatively calculate the exact probability, resulting in a Poisson-distributed variable, scaled by *p*_*ij*_. We chose our explicit definition to clearly separate the ecological and anthropological determinants of the true and observed number of reassortants, respectively. The model script can be found at https://github.com/fLuLab/AIV2344bEpizooticReassortment/blob/main/scripts/numberofreassortantsmodel/nreassortantsfinal.stan.

### A.2 Reassortant Class

Let *C* = {minor, moderate, major} be the ordered set of all possible reassortant classes. Each reassortant, *i* is assigned a class *y*_*i*_ = *c* ∈ *C*. To calculate the probability that a reassortant is assigned a particular class, we assume *y*_*i*_ follows a cumulative distribution in which minor *<* moderate *<* major [71]. We assume *y*_*i*_ originates from the categorisation of a latent (unobserved) continuous variable, 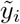, via threshold parameters, *τ*, that partition the latent scale into intervals corresponding to each class. Specifically, we define *τ*_minor−1_ = −∞ and *τ*_moderate+1_ = ∞:

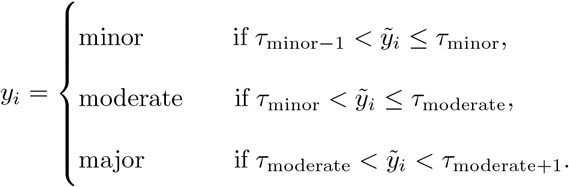

We modelled the latent variable 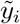 as the sum of four linear predictors, *η*_*i*_: i) the class identity of the reassortant immediately ancestral (with respect to HA) to reassortant *i*, ii) origin continent, iii) the number of segments changed relative to the immediately ancestral reassortant, and iv) the time interval between the MRCA of reassortant *i* and that for the most recent major reassortant. In addition, we modelled a penalised thin plate regression spline to smooth the time interval between the MRCA of reassortant *i* and that for the most recent major reassortant [72]. Briefly, for each knot, *κ*_*k*_, where *k* = 1, 2, …, *K*, we computed the distance, 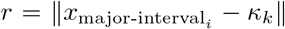, evaluated the radial basis function, *φ*(*r*) = *r*^2^ log *r*, and scaled the result by *γ*_*k*_. Altogether, we estimated *η*_*i*_ as:

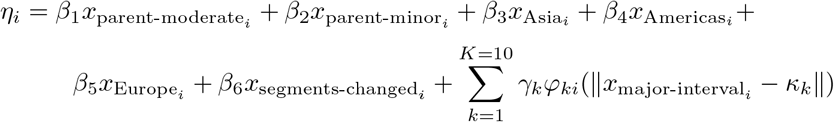

Let Φ be the cumulative distribution function of the Normal distribution. The probability that reassortant, *i*, is of class, *c*, is therefore given by:

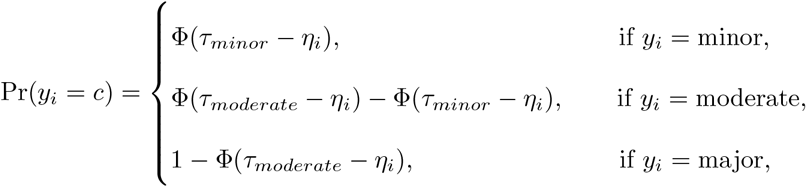

We specify the following weakly informative prior and hyperprior distributions for the reassortant class model:

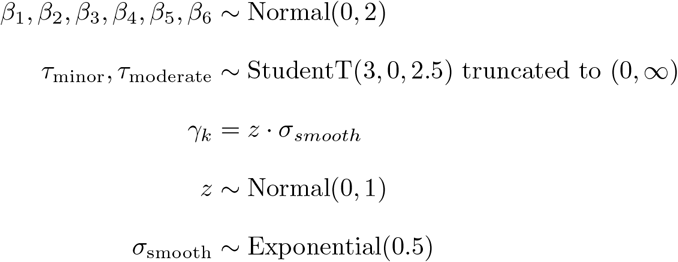

We fitted the model in Stan using BRMS v2.20.4. The model script can be found at https://github.com/fLuLab/AIV2344bEpizooticReassortment/blob/main/scripts/ordinalmodel/ordinalmodel.R.

### A.3 Diffusion Coefficient

For each reassortant, *i*, we estimated a weighted diffusion coefficient, *y*. We assume all *y*_*i*_ > 0 follow a Gamma distribution with shape, *κ*_*i*_, and rate, *θ*_*i*_, such that

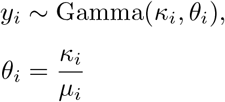

We assume shape parameter, *κ*_*i*_, is predicted as a log-linear function of the reassortant’s, indexed as 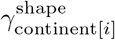 :

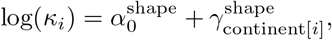

We model *µ*_*i*_ as the exponent of linear predictors for host richness, 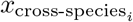; total persistence time, 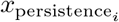; the number of sequences for each reassortant, 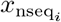; and the proportion of evolutionary time in wild Anseriformes spp, 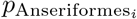, wild Charadriiformes spp, 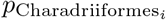, domestic Galliformes spp, 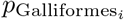 and other hosts, 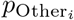. Because the proportions of evolutionary time spent in each host group are compositional (i.e., constrained to sum to one), we transform the proportions 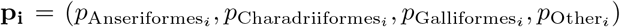, using an isometric log-ratio transformation [74]. Briefly, let **z**_*i*_ = ln(**p**_*i*_) denote the element-wise logarithm of the composition, and let **V** be a 4 × 3 orthonormal contrast matrix defining the sequential binary partitions: (i) Charadriiformes vs. Anseriformes, (ii) (Anseriformes + Charadri-iformes) vs. Galliformes, and (iii) (Anseriformes + Galliformes + Charadriiformes) vs. Other hosts. Because log-ratio transformations require strictly positive components, we substituted zero-valued proportions with a small positive constant, *δ*, and rescaled. The log-ratios, **x**_ilr,*i*_, are given by:

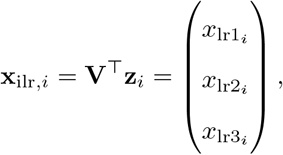

where

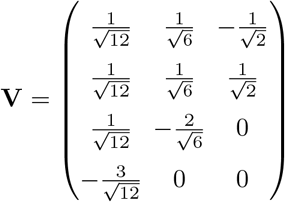

We included zero-centred random intercepts for the date of the reassortant MRCA grouped by calendar year, *γ*_year[*i*]_; and reassortant origin continent. We allowed the group-level effect of origin continent to vary across uncorrelated slopes of the relative proportions of evolutionary time in wild Anseriformes spp/Charadriiformes, 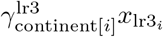 and (Anseriformes + Charadriiformes) vs. Galliformes, 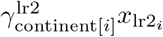:

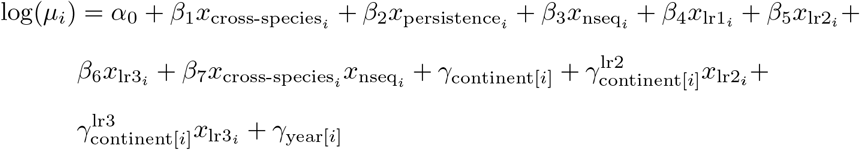

Finally, we specify the following prior and hyperprior distributions for the diffusion coefficient model:

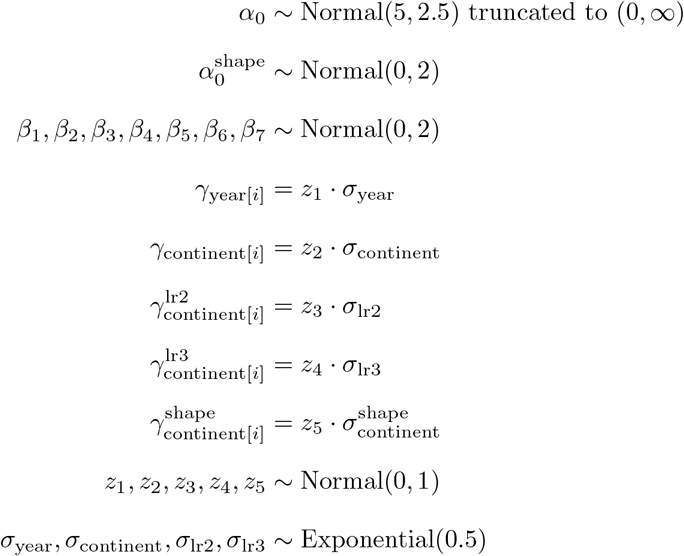

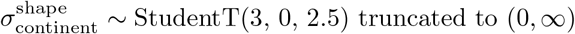

We fitted the model in Stan using BRMS v2.20.4. The model script can be found at https://github.com/fLuLab/AIV2344bEpizooticReassortment/blob/main/scripts/diffusionmodel/diffusionmodel.R.

## APPENDIX B Supplementary Figures

**Fig. S1.**
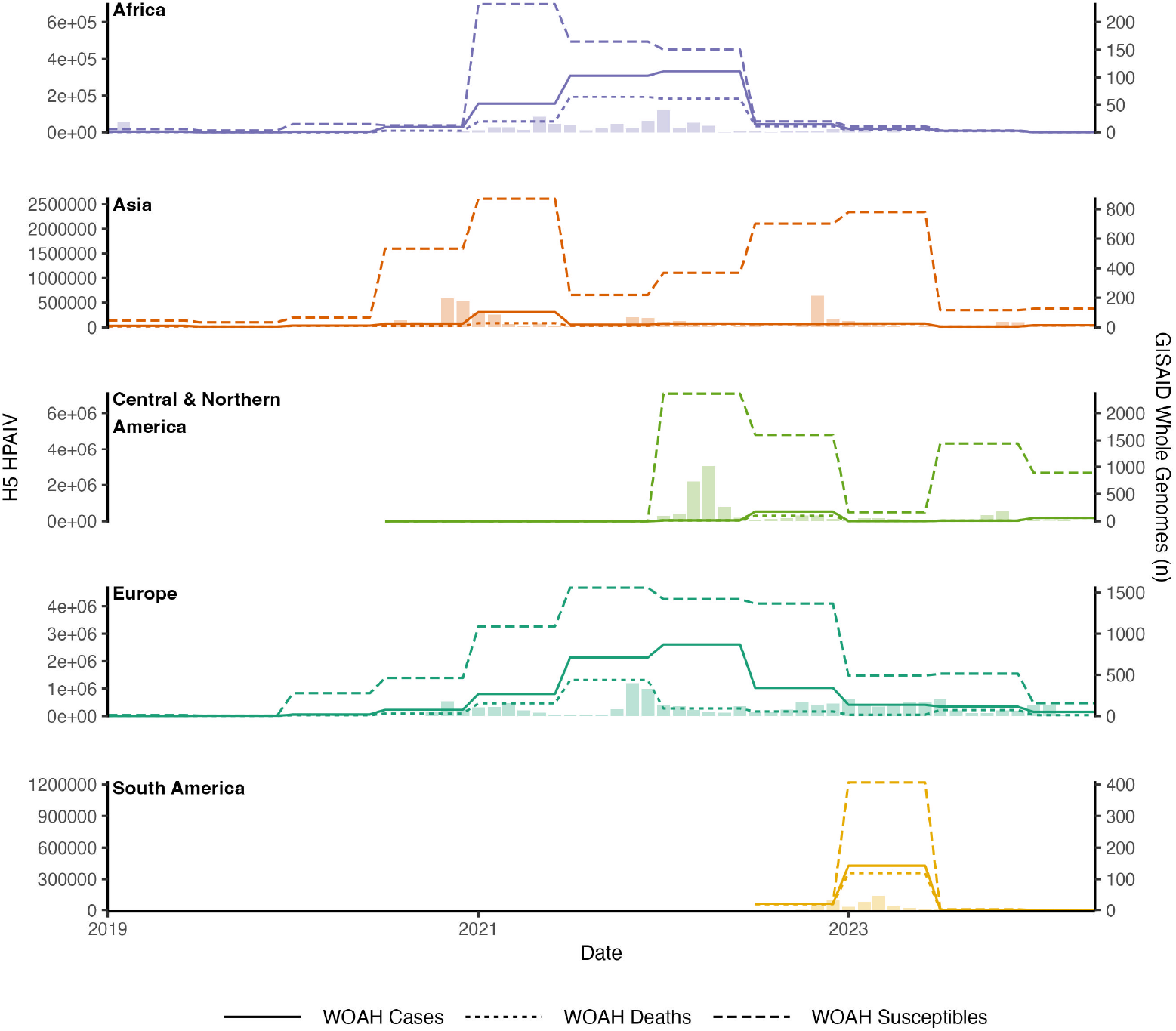
High Pathogenicity H5 reports from World Organisation for Animal Health. We downloaded all biannual reports concerning high pathogenicity H5 avian influenza between January 2019 and July 2024. Stratified by reporting continent, we compare the number of reported cases (infected or sick animals and animals that died from the disease), deaths (animals that died from the disease) and susceptibles (animals present in the outbreak at the beginning of the biannual reporting period), relative to the number of GISAID whole genome sequences. In our statistical models, we selected the number of susceptibles to represent H5 HPAIV incidence due to inconsistencies in deaths and cases (e.g deaths sometimes exceed the number of cases). Bars correspond to the number of whole genomes and their sampling date

**Fig. S2.**
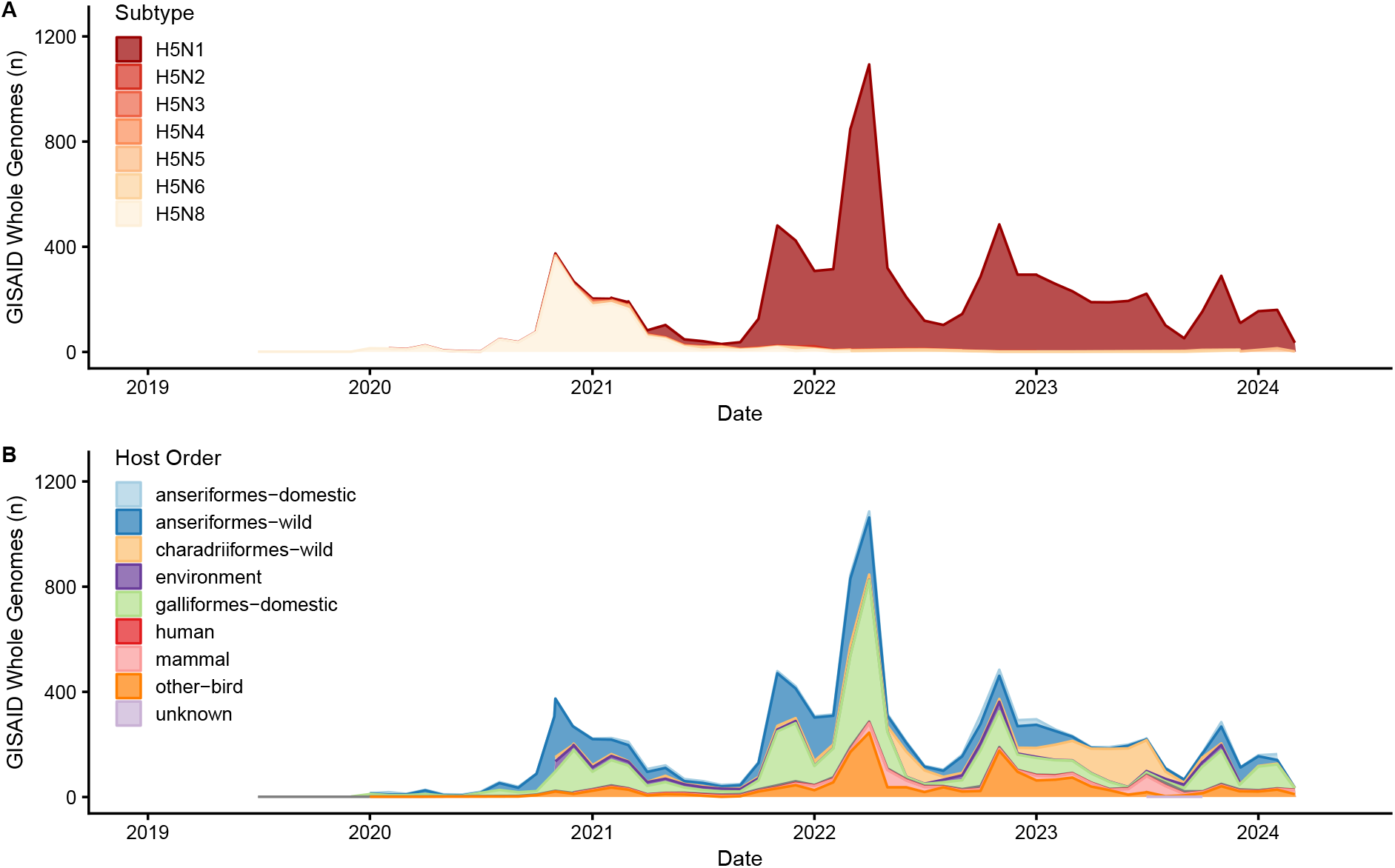
Epidemic transitions throughout 2020-2024 2.3.4.4b Panzootic. Our data encompasses key changes in the epidemiology of H5 high pathogenicity avian influenza virus. In particular, these data include distinct changes in host composition (A) and influenza subtype (B) as the panzootic progressed

**Fig. S3.**
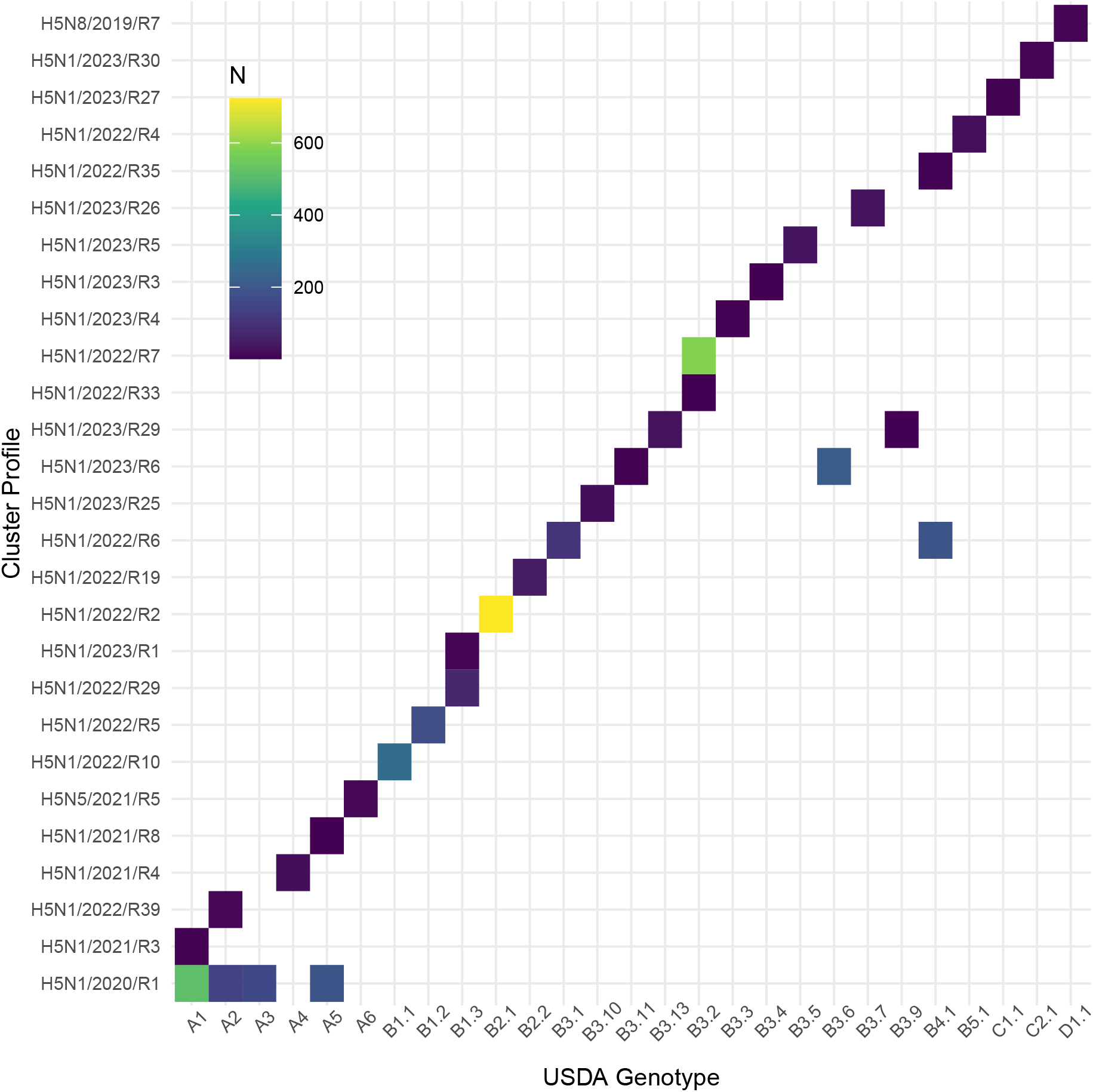
Comparison between USDA genoflu (https://github.com/USDA-VS/GenoFLU) and the nomenclature used in this study. From 3,648 isolates for which both nomenclature were available, our reasssortant profiles demonstrate consistency with those assigned by genoflu.

**Fig. S4.**
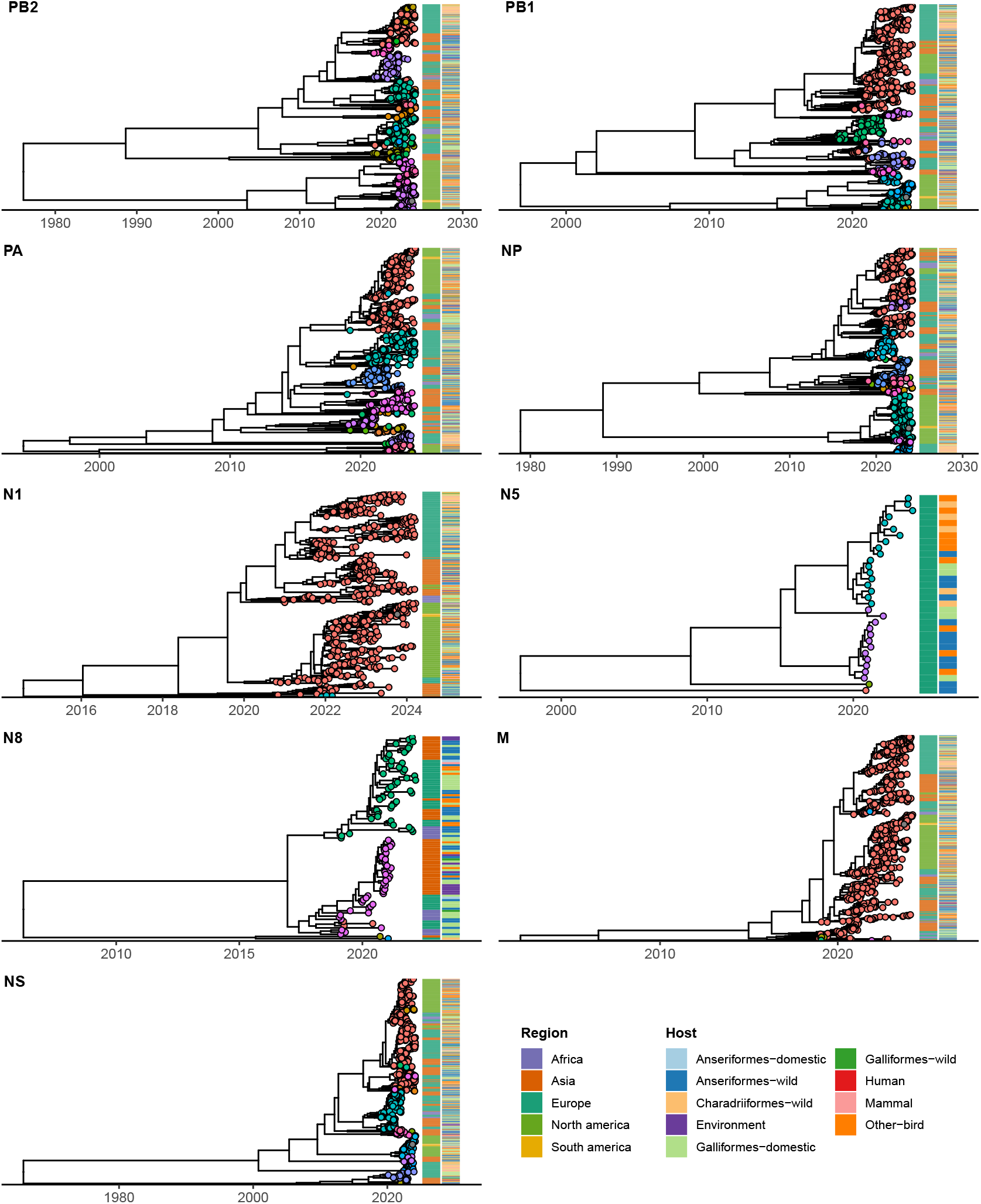
Individual segment time-resolved phylogenies (excluding HA), annotated by reassortant cluster. In this study, we defined reassortants using hierarchical clustering Specifically, we clustered all whole genomes in our dataset by pairwise nucleotide distances, setting a 0.5% threshold for the longest three gene segments (PB2, PB1 and PA) and a 1% threshold for the remaining segments. The tips of each segment phylogeny are coloured by their assigned cluster. Adjacent to each phylogeny, the first column is coloured by sampling region and the second by host.

**Fig. S5.**
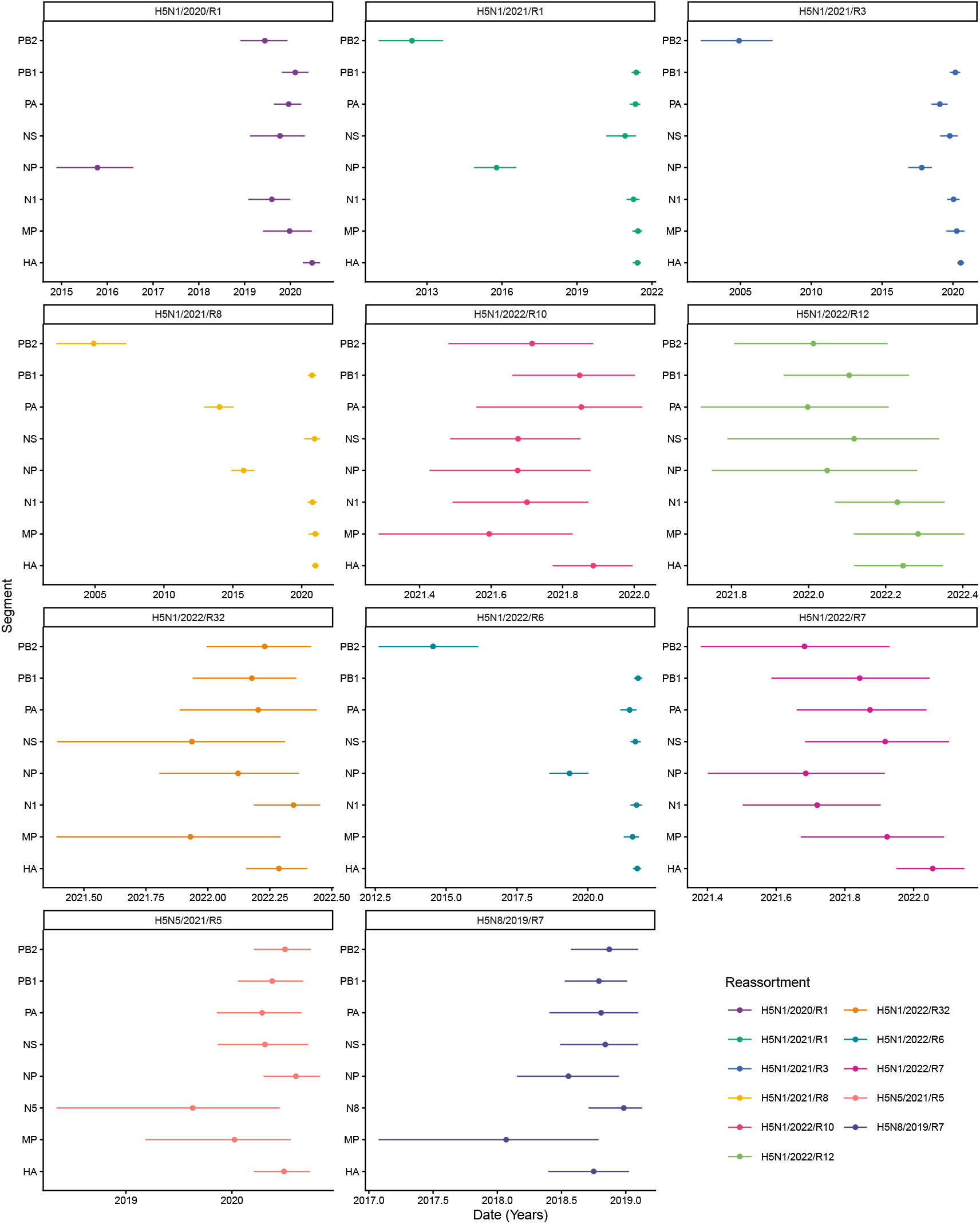
Comparison between segments of the date of the most recent common ancestor (MRCA) for the eleven most frequently sampled reassortants. Estimated from Bayesian time-resolved phylogenies, each point reresents the median posterior estimate of the date of the MRCA, while lines show the 95% highest posterior density.

**Fig. S6.**
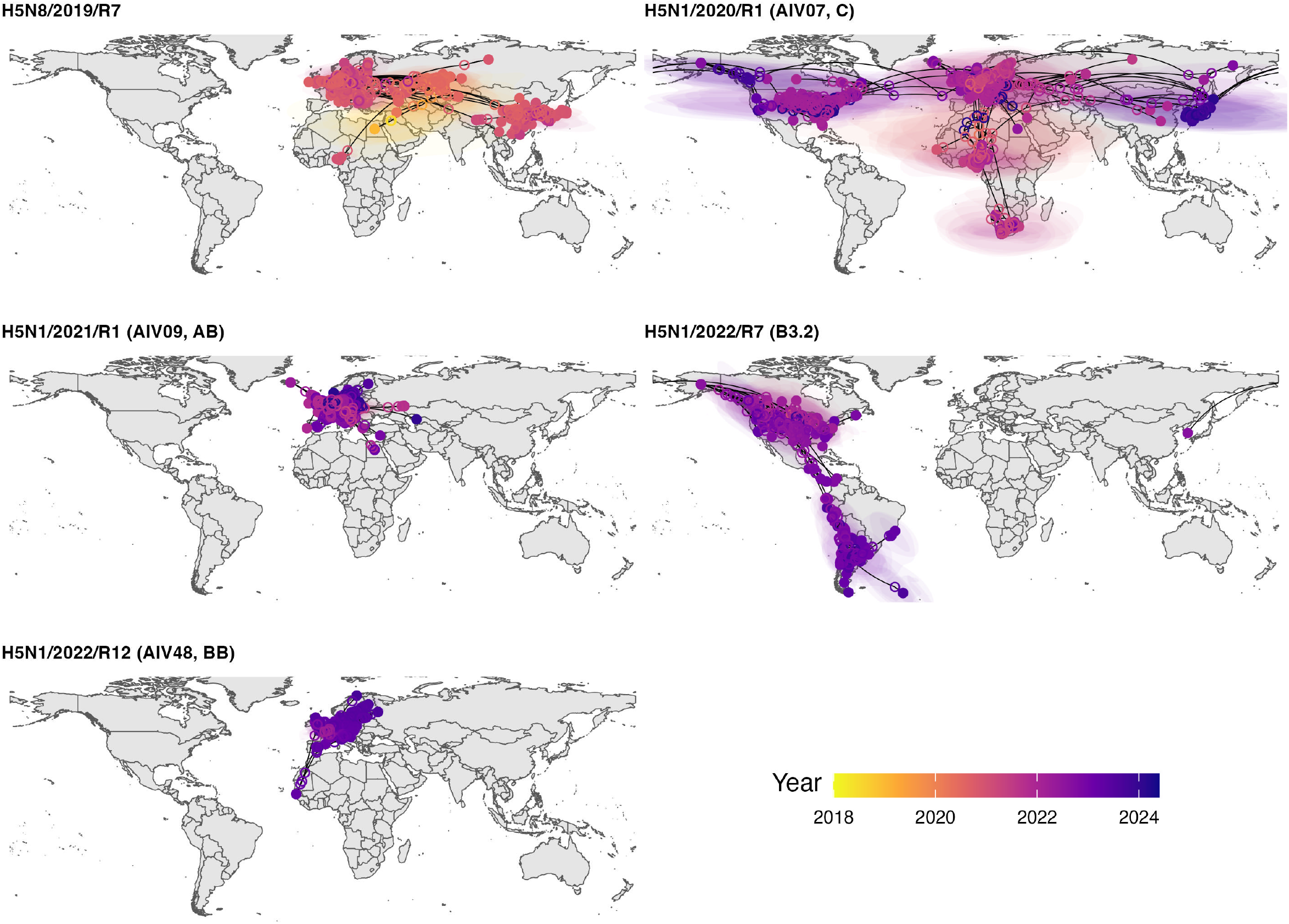
Continuous Phylogeography for Major Reassortants. For each reassortant, we inferred global circulation patterns throughout time. Solid dots are the location and time of tip sequences, while hollow dots correspond to the estimated location of intermediate nodes. Shaded polygons show the 95% highest posterior density estimated for each internal node.

**Fig. S7.**
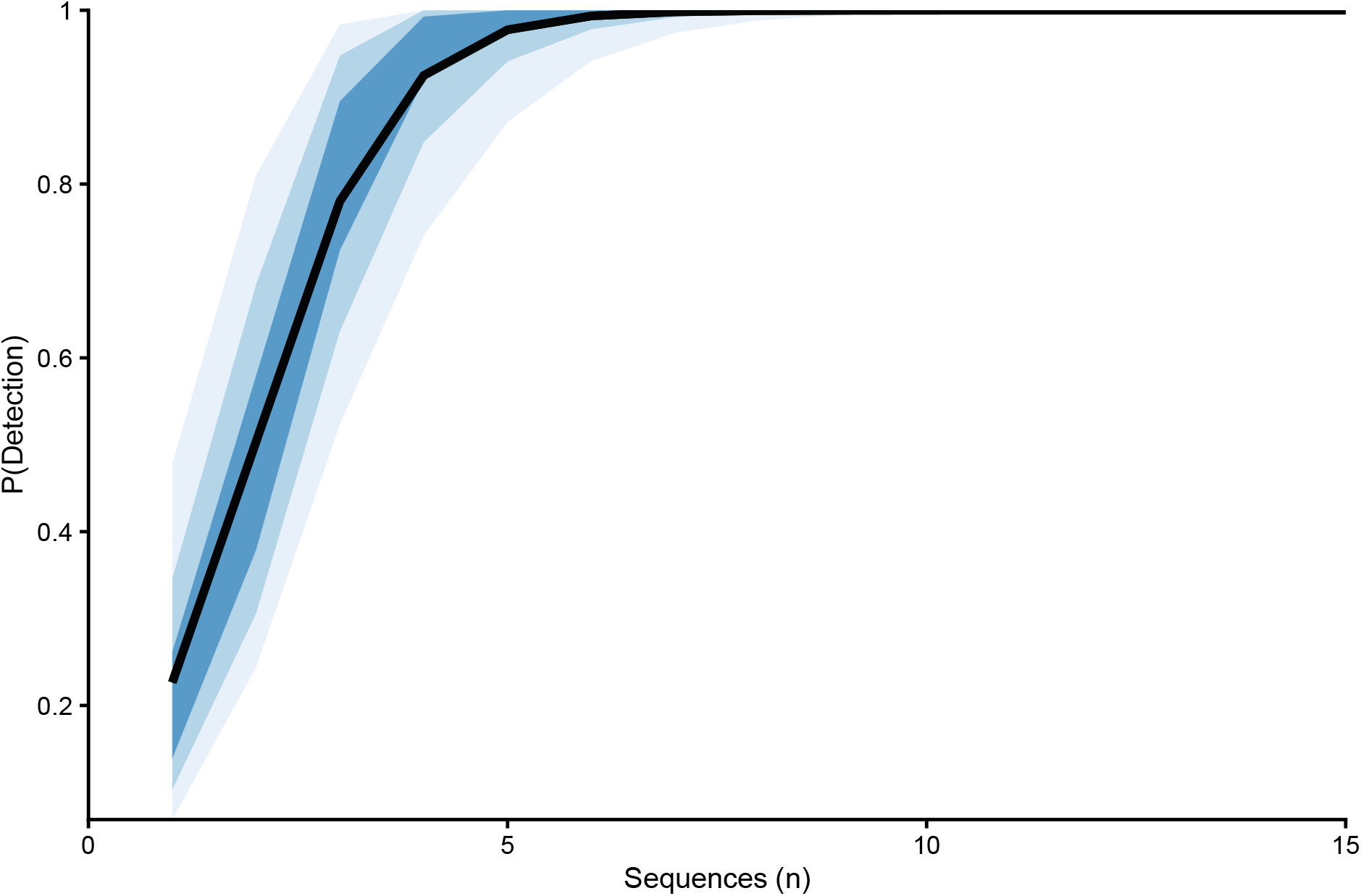
Conditional effect of the number of sequences per month on the probability that a latent reassortant in the environment is eventually detected. Shaded regions indicate highest posterior density intervals for each estimate, corresponding to the 95%, 85%, and 50% levels from light to dark.

**Fig. S8.**
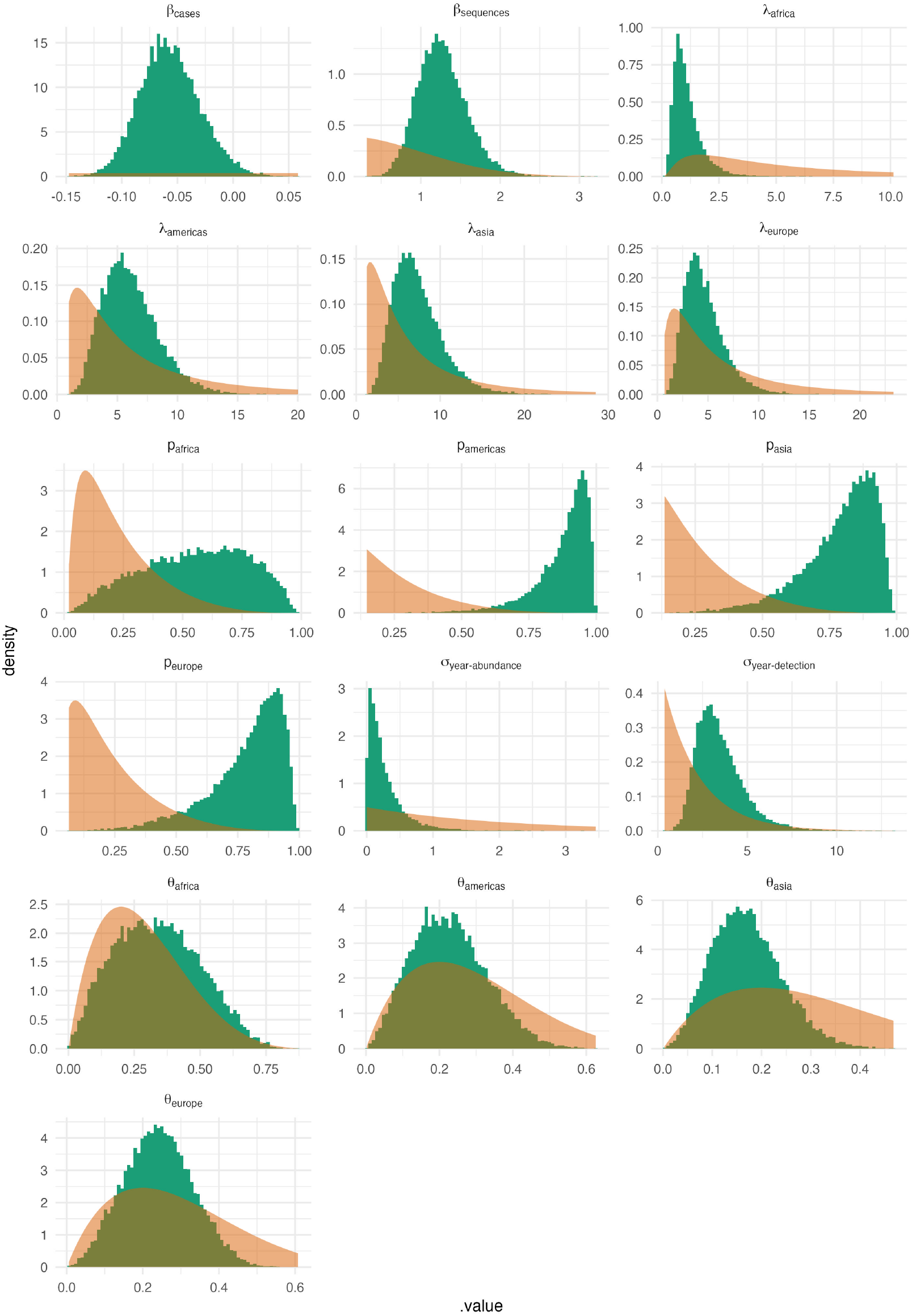
Prior (orange) and posterior distributions (green) for select parameters of the ‘numbers of reassortant’ model. In most cases the two distributions show minimal overlap, which indicates model ‘learning’ from the data. For *θ* parameters, there is a close overlap between the some posterior and prior distributions, indicating these parameters are only weakly identifiable

**Fig. S9.**
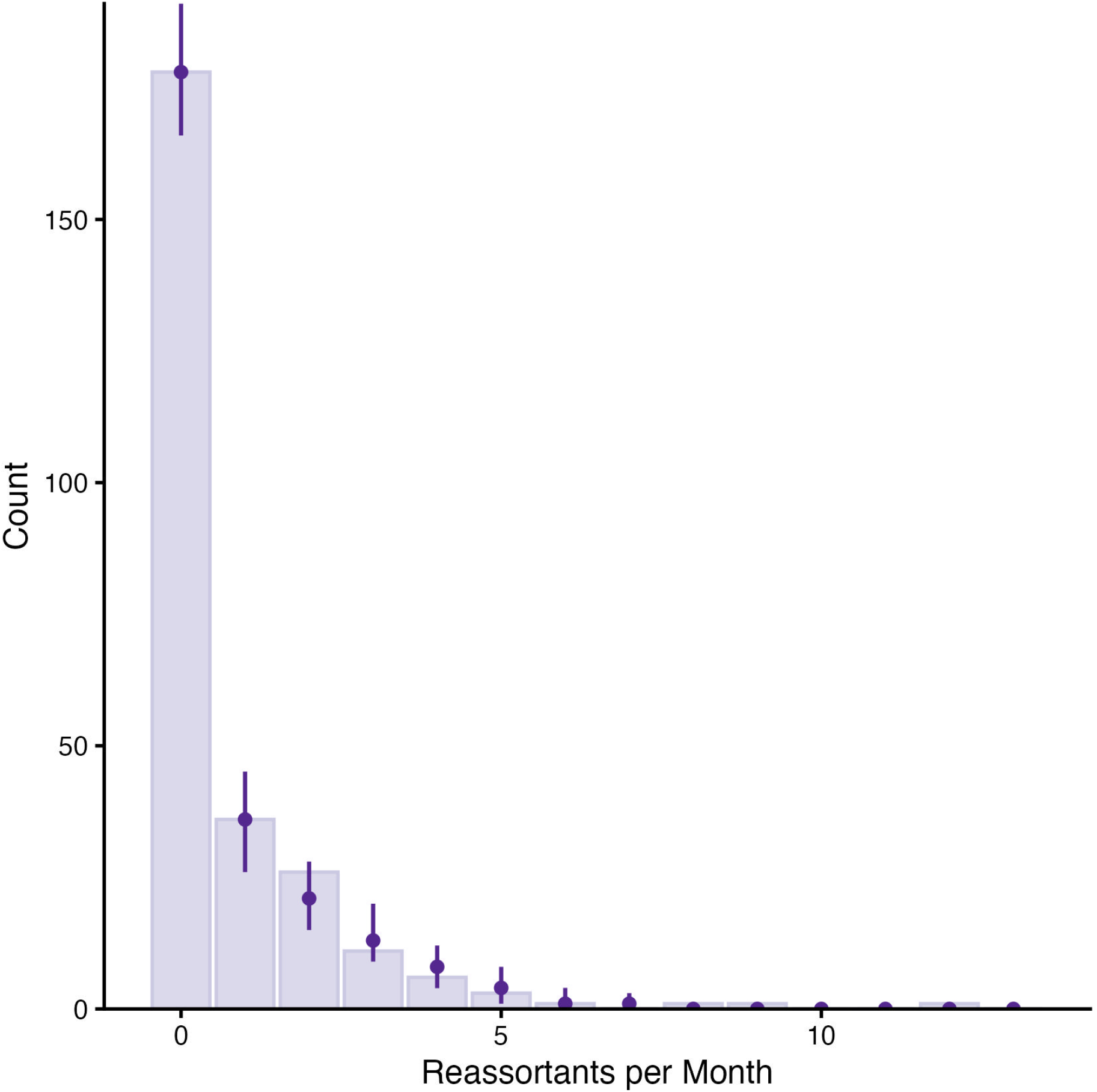
Posterior predictive check for the ‘numbers of reassortant’ model. The underlying bar plot represents the empirical data distribution. Each point corresponds to the median estimate of the posterior predictive distribution, with the associated error bars displaying the 95% Highest Posterior Densities (HPD).

**Fig. S10.**
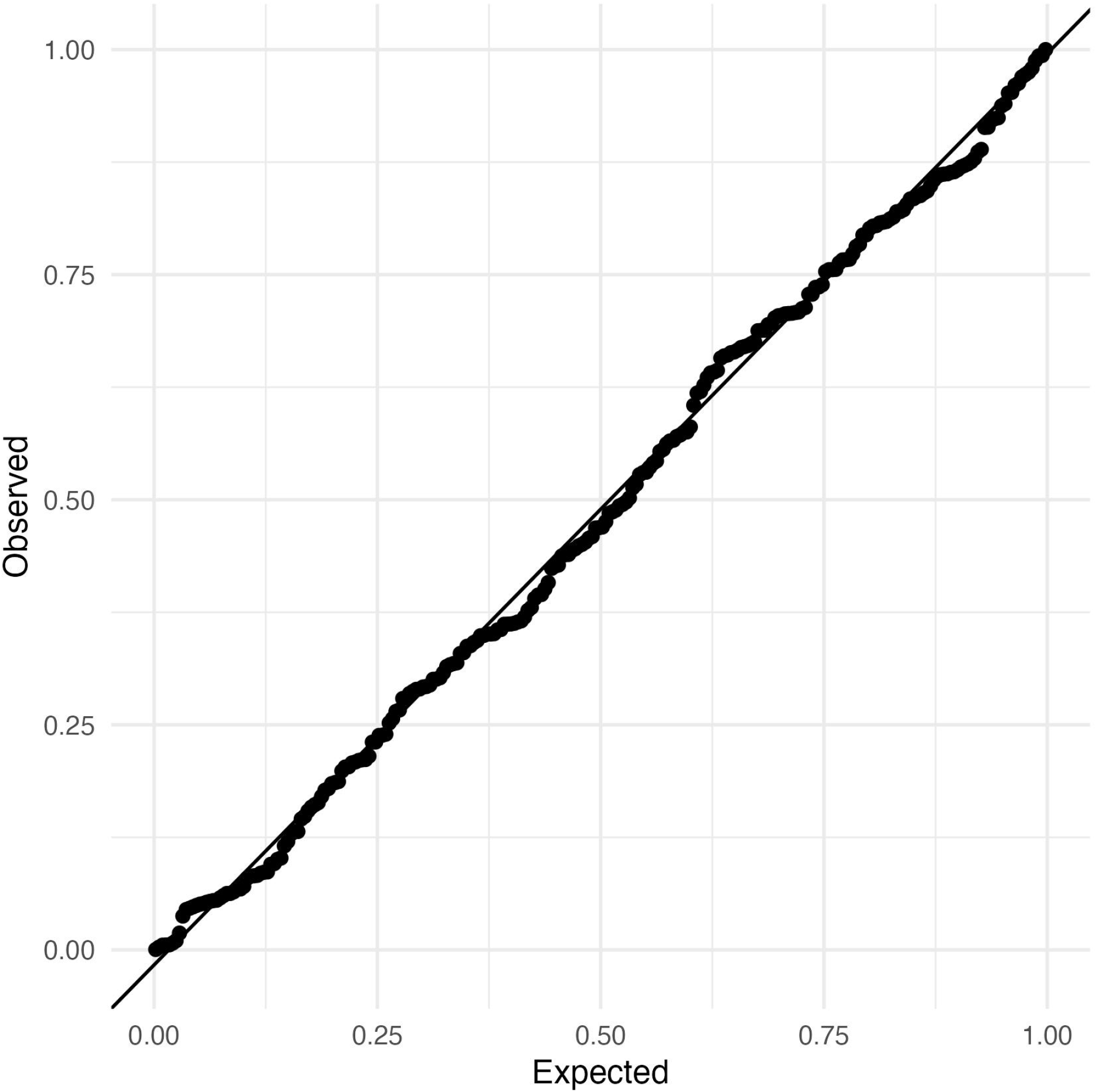
Quantile-Quantile residual diagnostic plot for the numbers model. Scaled residuals were calculated using DHARMa [118]

**Fig. S11.**
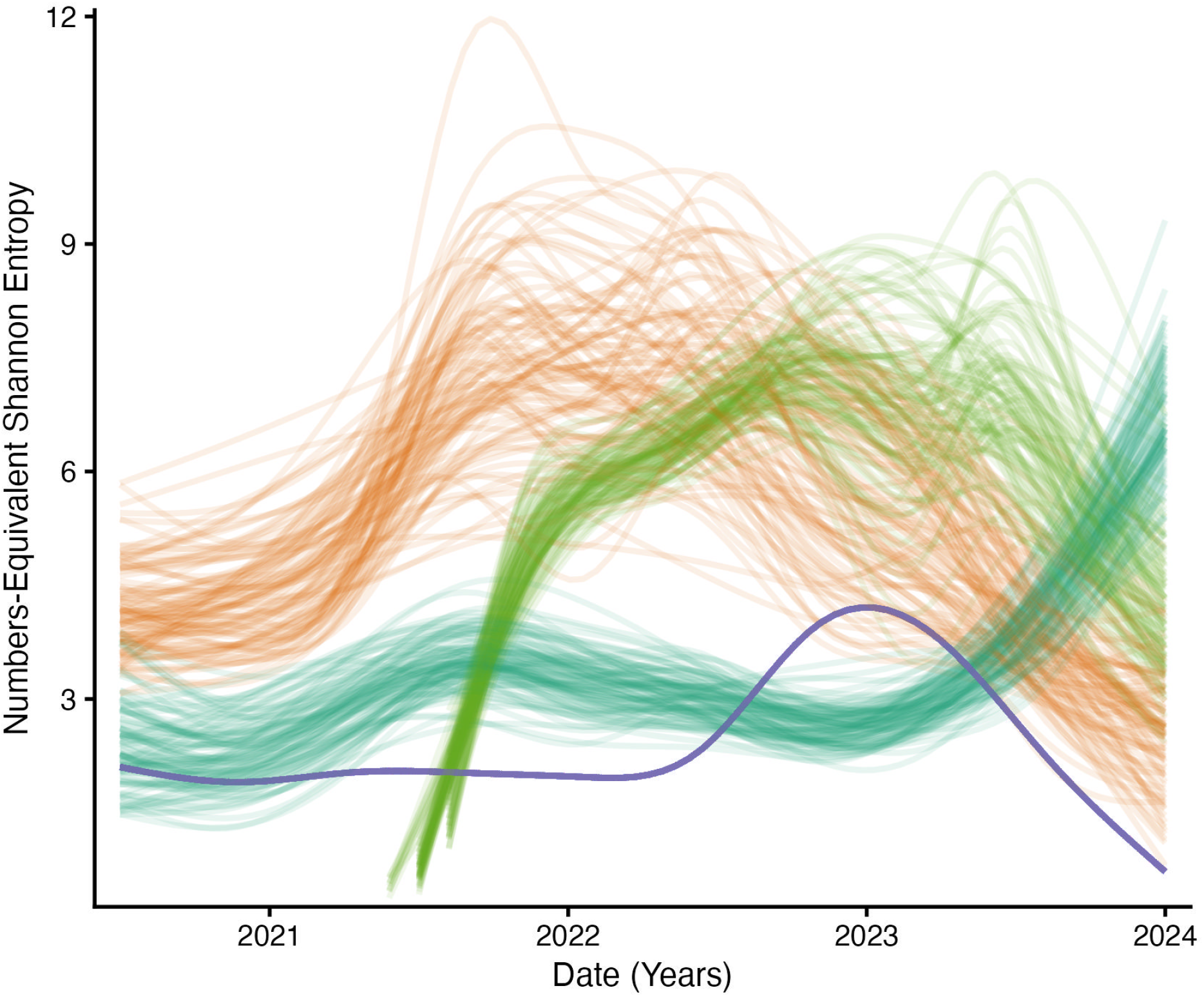
Downsampling of Numbers-Equivalent Shannon Entropy. In our dataset, Africa was the least frequently sampled continent. To evaluate the robustness of numbers-equivalent Shannon entropy to uneven sampling, we reanalysed the data while holding the number of whole genomes analysed per continent equal to the number sampled from Africa. For 100 iterations, we repeatedly resampled genomes from all continents. Each line in the plot represents a single resample per continent. Overall, the patterns within each continent remained broadly consistent across the resampled data relative to the full dataset

**Fig. S12.**
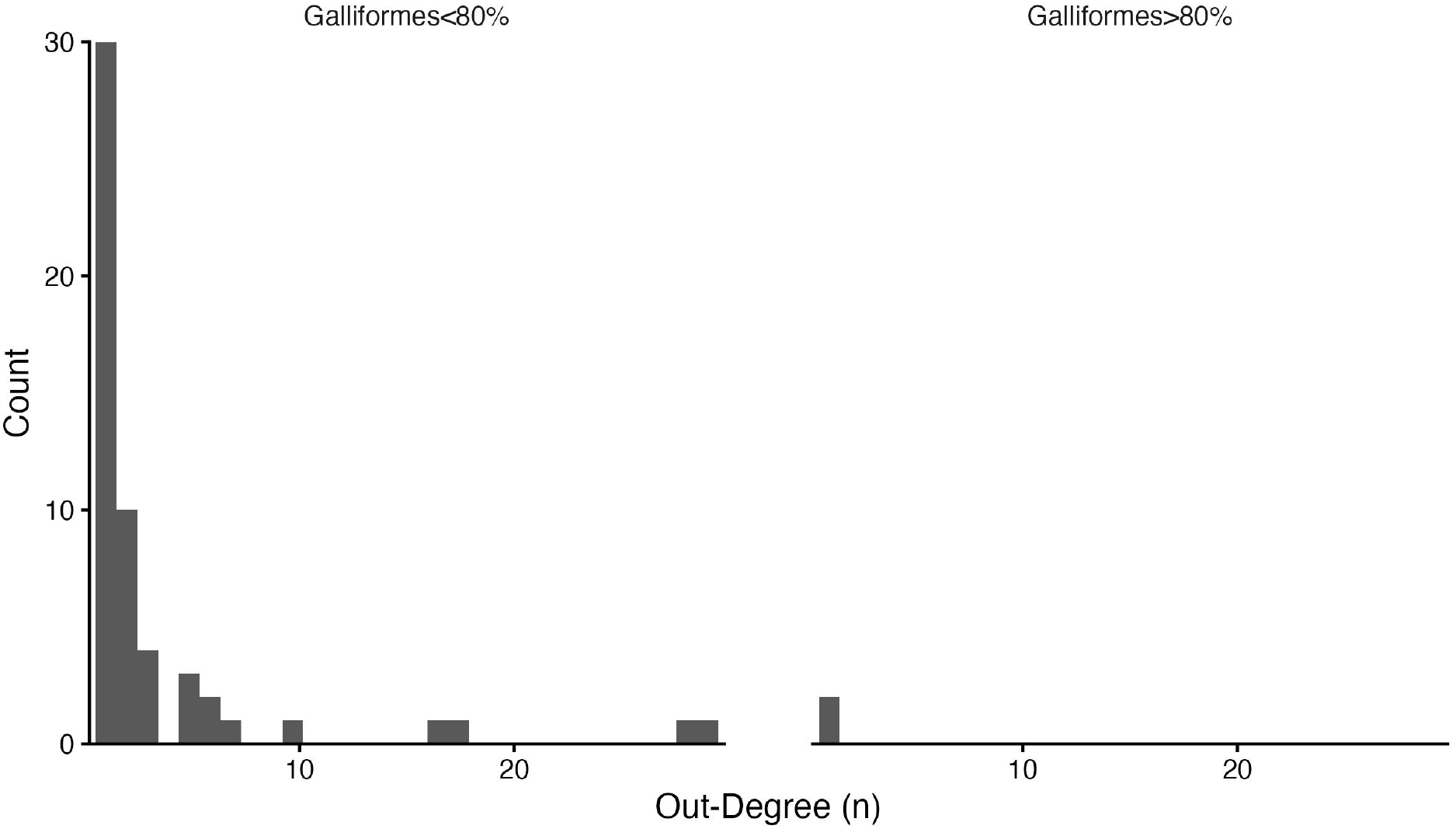
Number of Offspring Reassortants in Galliformes. In our dataset, we observed that few reassortants originated in Galliformes. To investigate this further, we compared the ‘out-degree’ distribution (i.e. the number of offspring reassortants) between reassortants with greater than or equal to 80% sequences isolated from Galliformes spp, and those with less than 80%. We found that reassortants with at least 80% of sequences sampled from Galliformes were significantly less likely to be ‘parent’ reassortants.

**Fig. S13.**
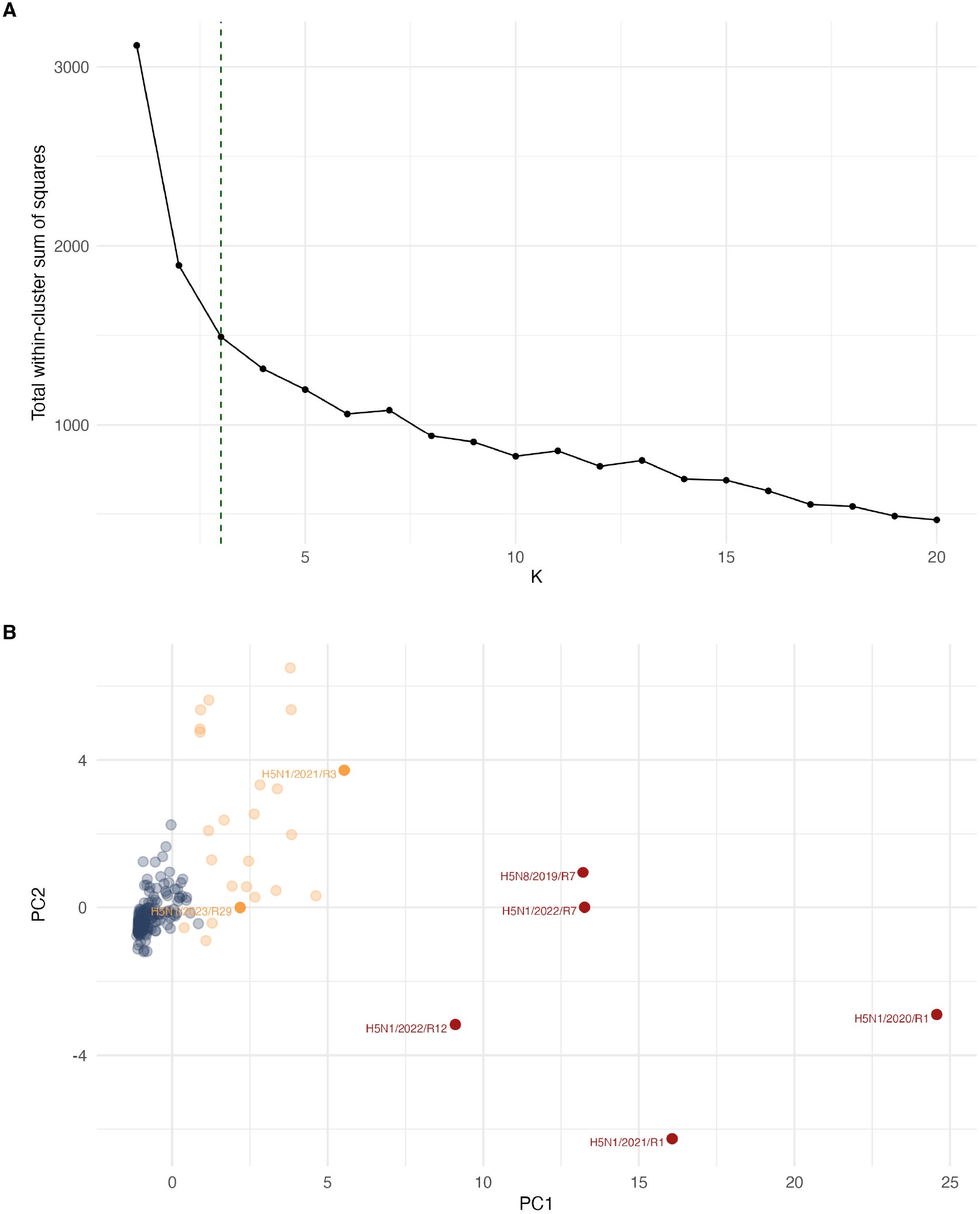
K-Means Clustering of Reassortant Phylodynamic Profiles. A) Selection of k = 3 as optimal number of clusters for further analysis. B) Two dimensional representation of our phylodynamic profiles, where each point represents a single reassortant and is coloured by assigned cluster: major (red), moderate (yellow), and minor (blue).

**Fig. S14.**
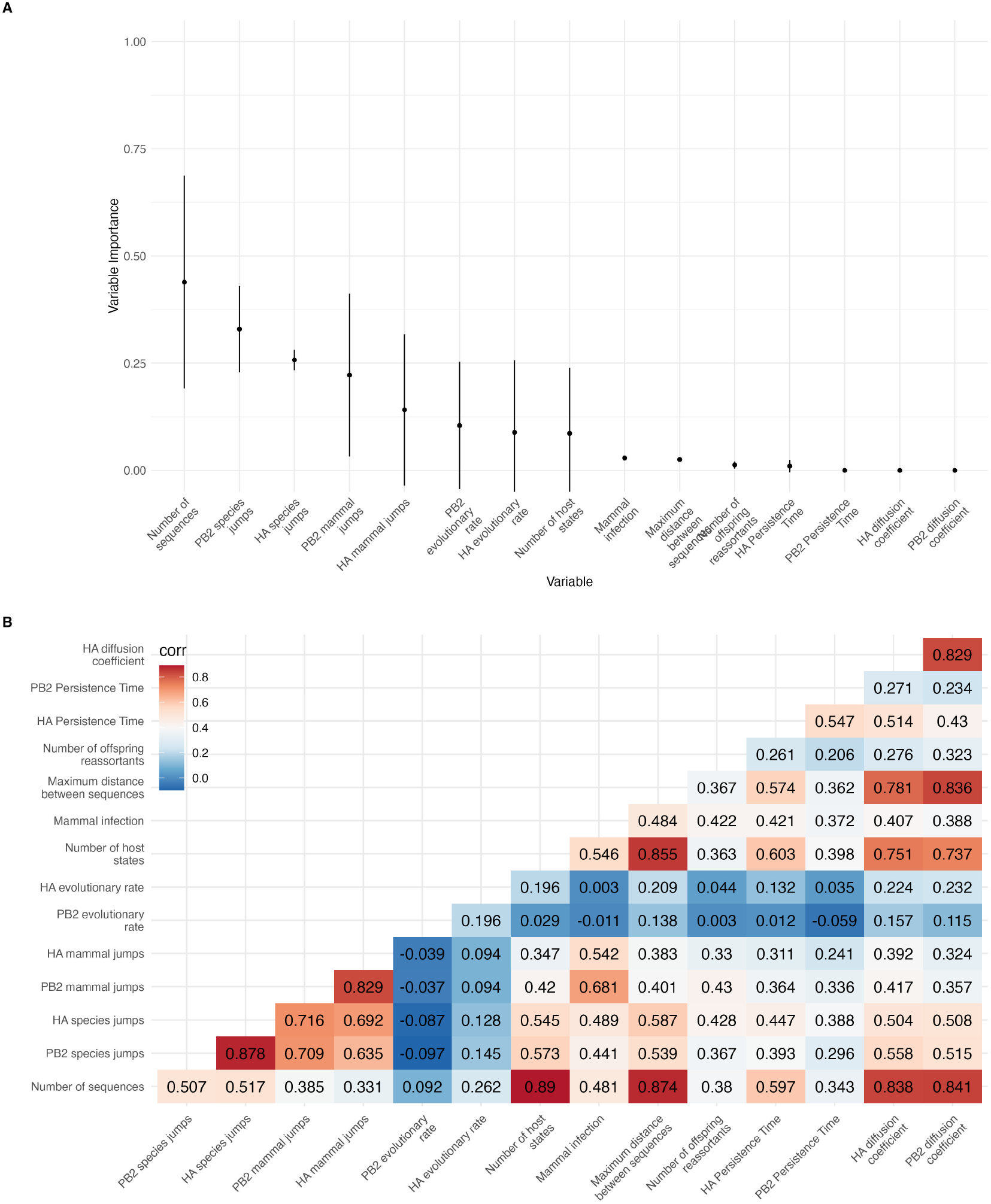
K-Means Clustering of Reassortant Phylodynamic Profiles. A) Relative influence of phylodynamic variables on cluster allocation. We iteratively exclude one variable and calculate the similarity in clustering patterns between the original and iterative result using the Adjusted Rand Index (ARI). We define variable importance as 1-ARI. B) Spearman Correlation coefficients calculated between transformed phylodynamic variables. Metrics counting the number of species jumps, the number of mammal jumps and the diffusion coefficient typically show a strong (¿0.7) positive correlation between HA and PB2.

**Fig. S15.**
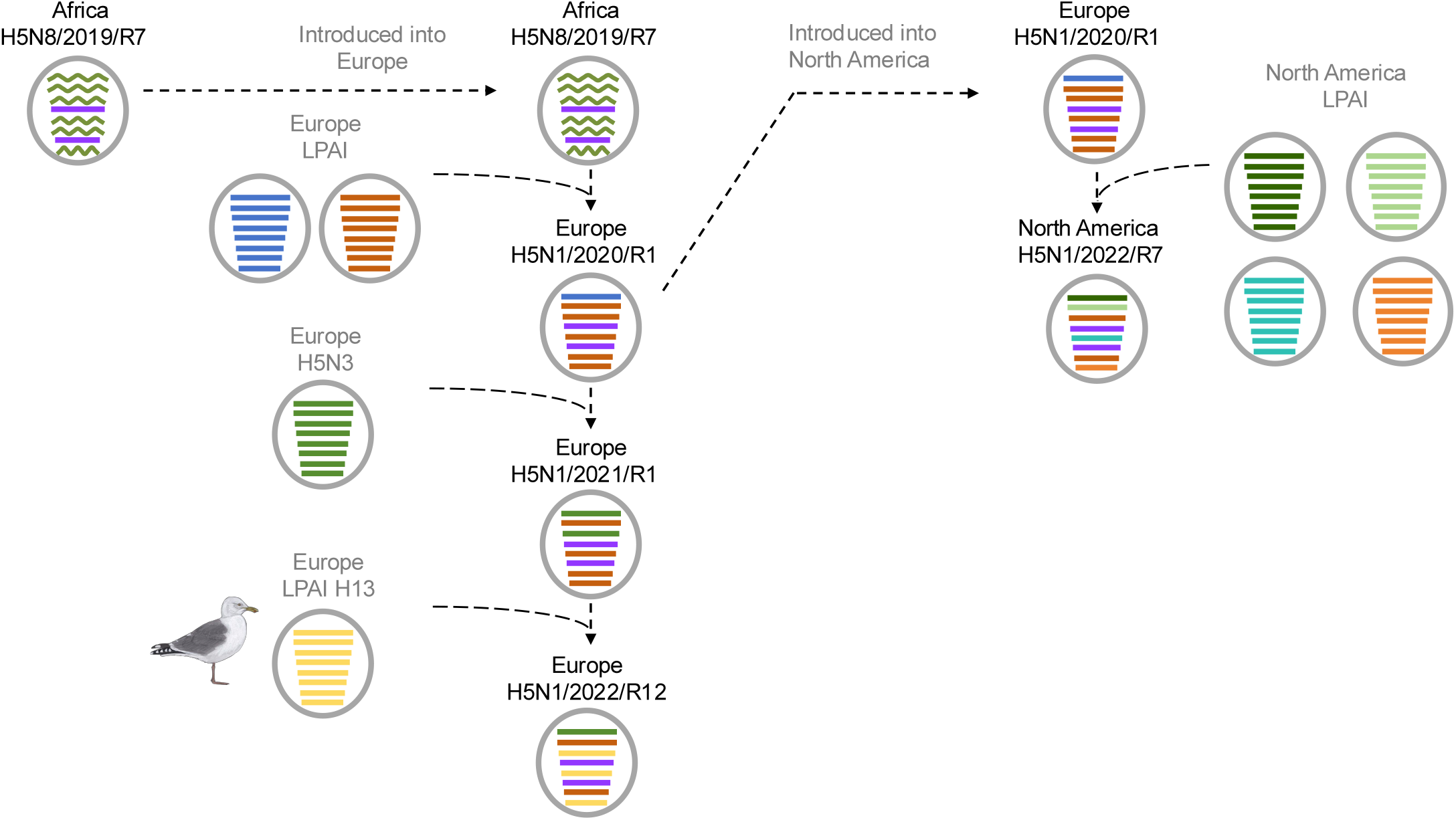
Progression of Major reassortants across the 2020-2024 2.3.4.4b H5 panzootic

**Fig. S16.**
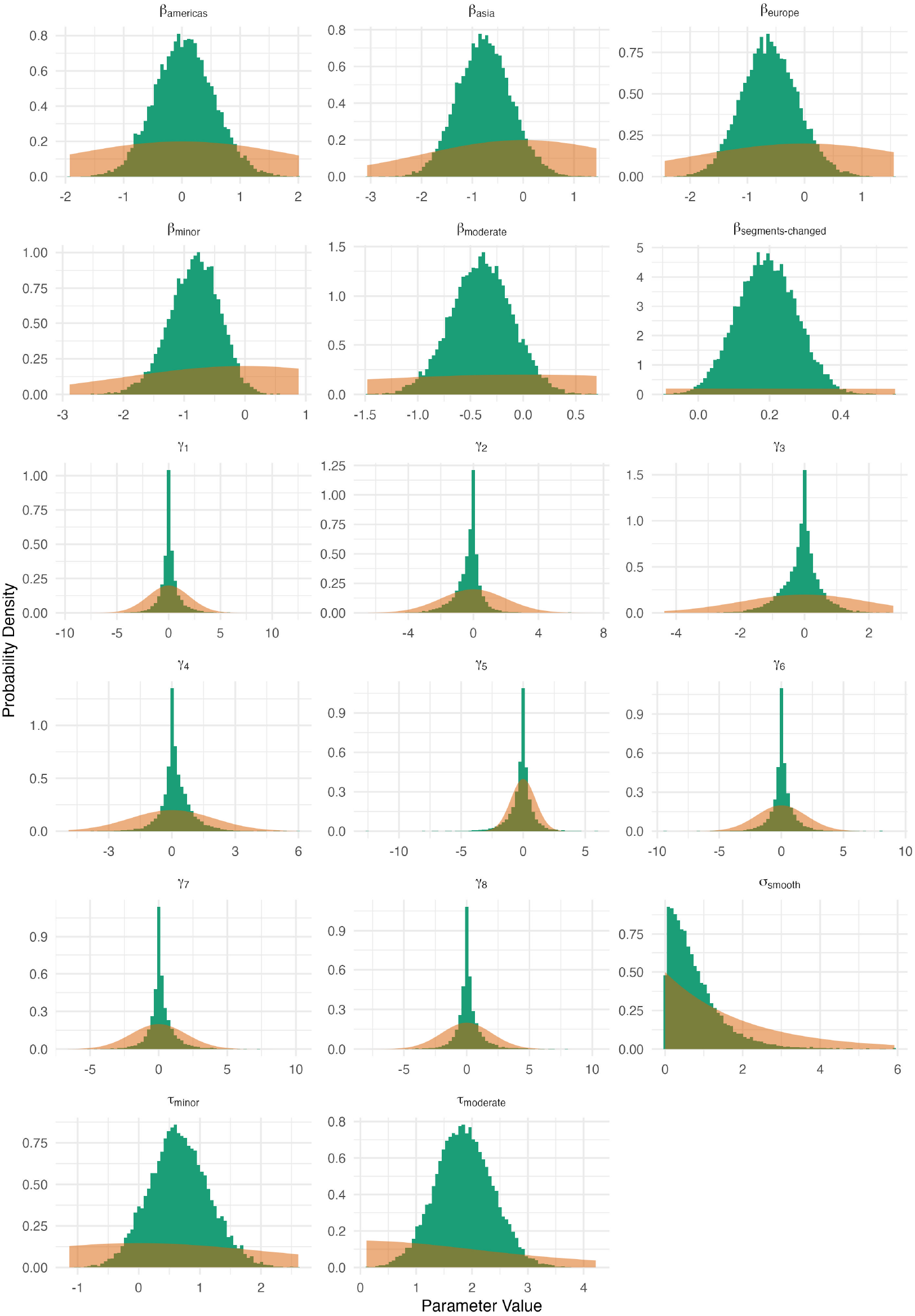
Prior (orange) and posterior distributions (green) for select parameters of the reassortant class model. In all cases the two distributions show minimal overlap, which suggests the posterior distributions are well-informed by the model likelihood.

**Fig. S17.**
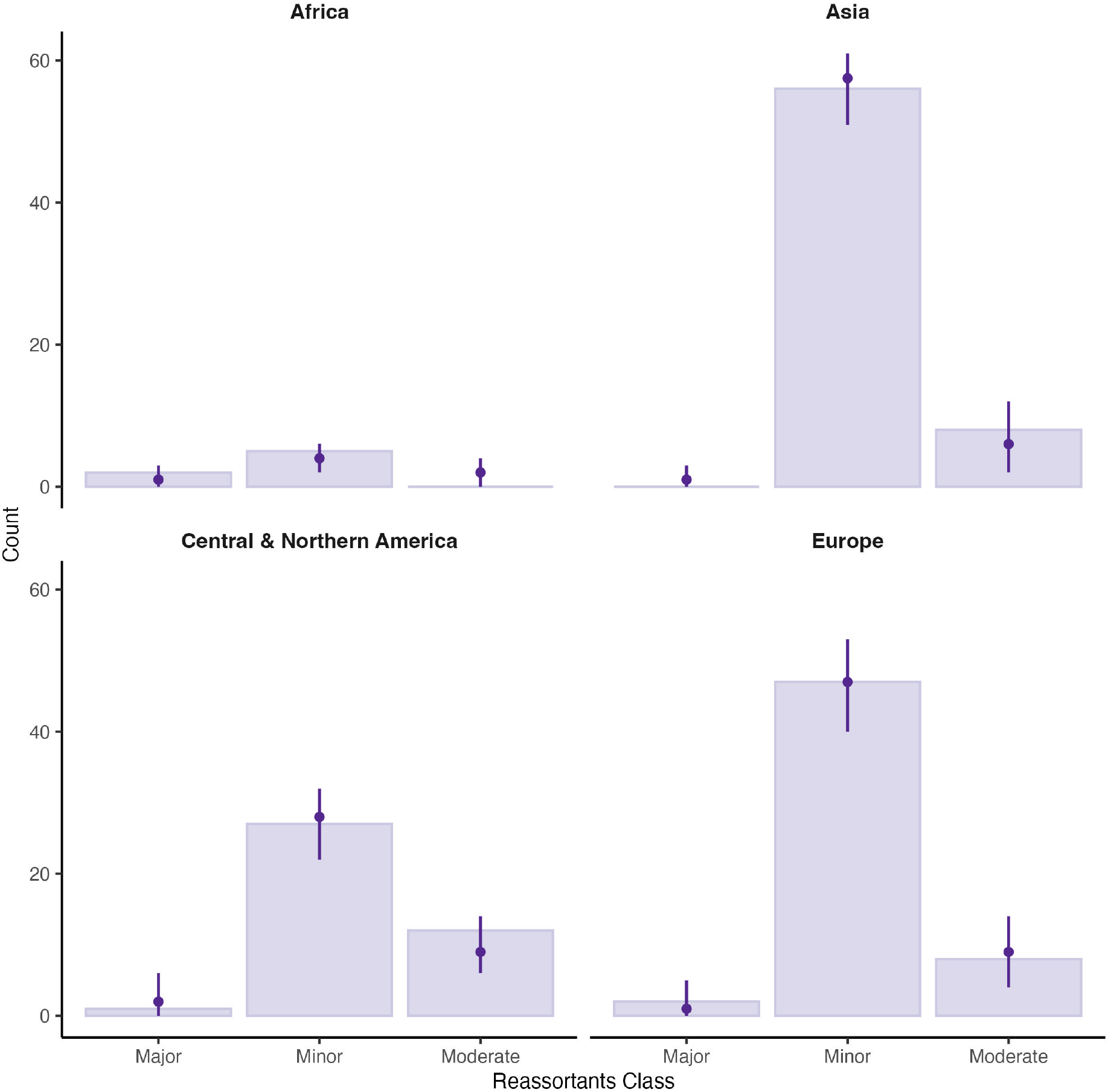
Posterior predictive check for reassortant class model. Stratified by continent, each underlying bar plot represents the empirical data distribution. Each point corresponds to the median estimate of the posterior predictive distribution, with the associated error bars displaying the 95% Highest Posterior Densities (HPD).

**Fig. S18.**
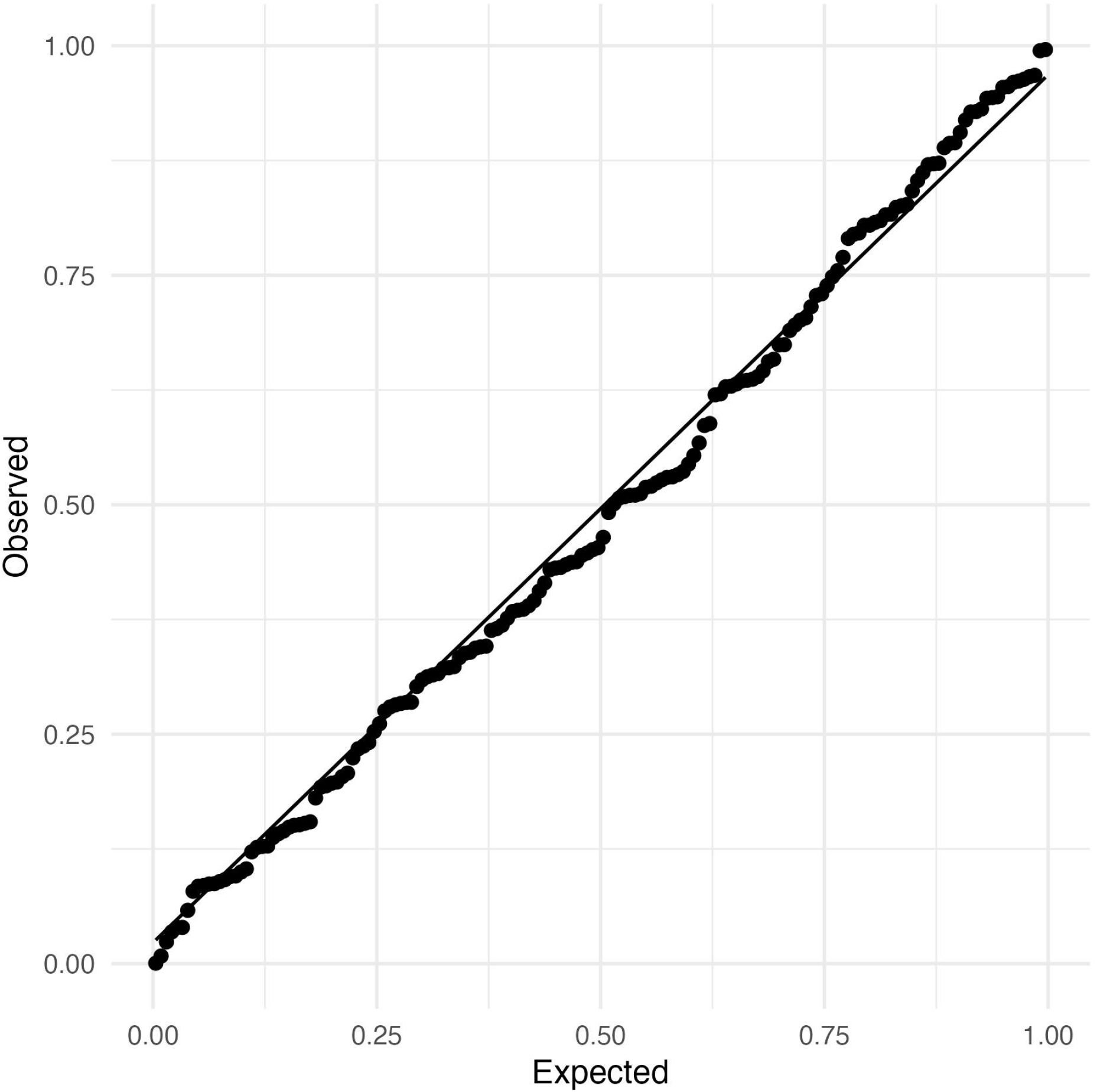
Quantile-Quantile residual diagnostic plot for the reassortant class model. Scaled residuals were calculated using DHARMa [118]

**Fig. S19.**
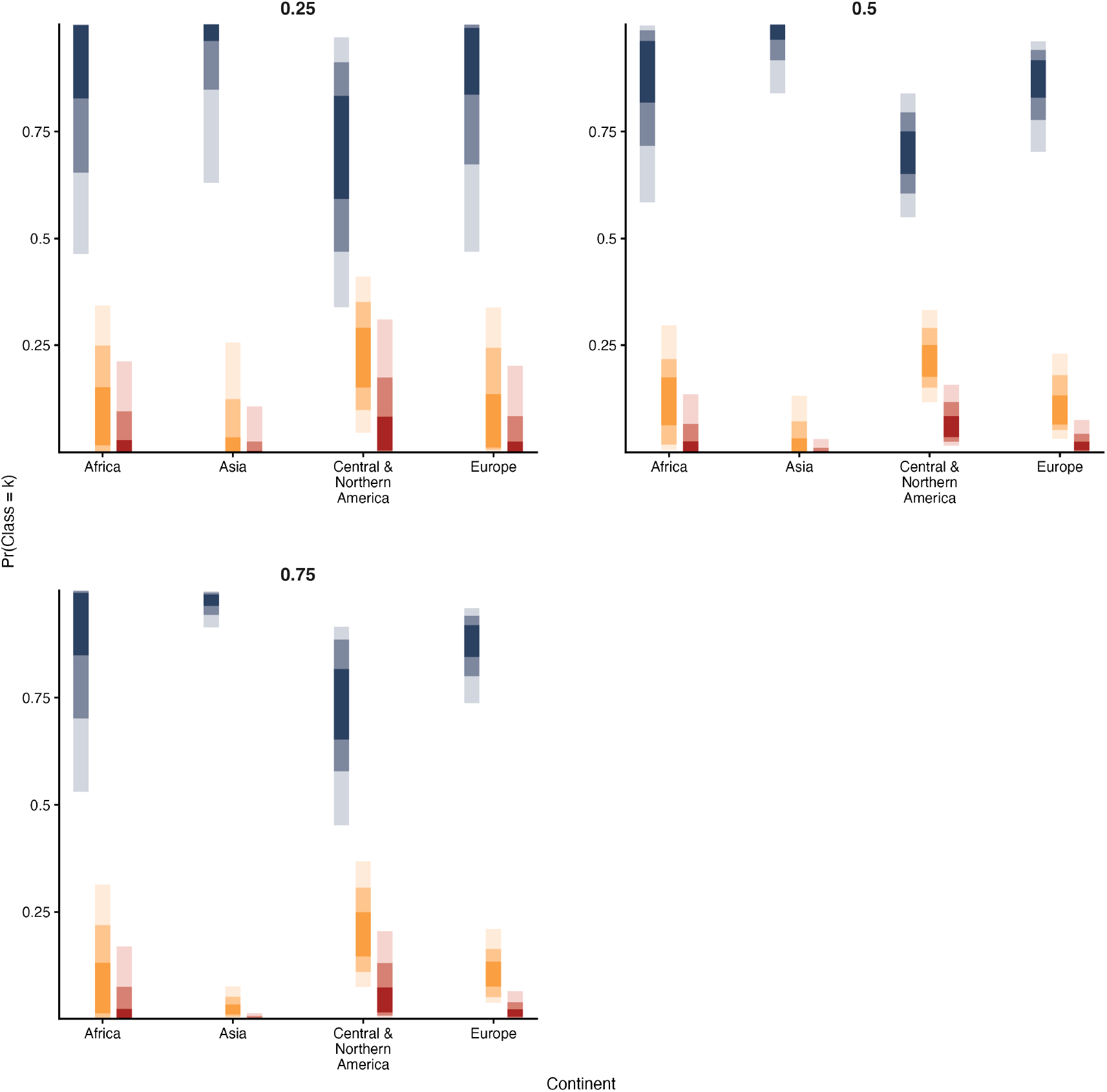
Across all NFLG sequences in our data (n = 9,935), the proportion of wild sequences was greatest in Asia (p = 0.757), followed by Europe (p = 0.635), Central & North America (p = 0.555), South America (p = 0.513) and Africa (p = 0.4714). We considered whether the proportion of sequences sampled from wild birds might alter the predicted probabilities for each reassortant class, across regions. To achieve this, we extended our original ordinal model, assuming the effect of the proportion of wild sequences remains constant across regions. Averaged across the empirical data distribution, we would expect a 25% increase in the proportion of wild birds to decrease the probability that a given reassortant is ‘major’ by 0.00244 (95% HPD: -0.0311;0.0864).We then evaluated a counterfactual scenario where the proportion of wild bird sequences was fixed at 0.25, 0.5, and 0.75 for each region, assuming all else equal. At proportion of wild bird sequences of 0.25, 0.5, and 0.75, our results were qualitatively consistent with the main analysis. Specifically, major reassortants are more likely to have emerged from Central/North America at 25% (0.0791 (5.46e-03; 0.371)), 50% (0.0732 (2.05e-02; 0.173)) and 75% (0.0651 (9.86e-03; 0.239) wild sequences than Europe.

**Fig. S20.**
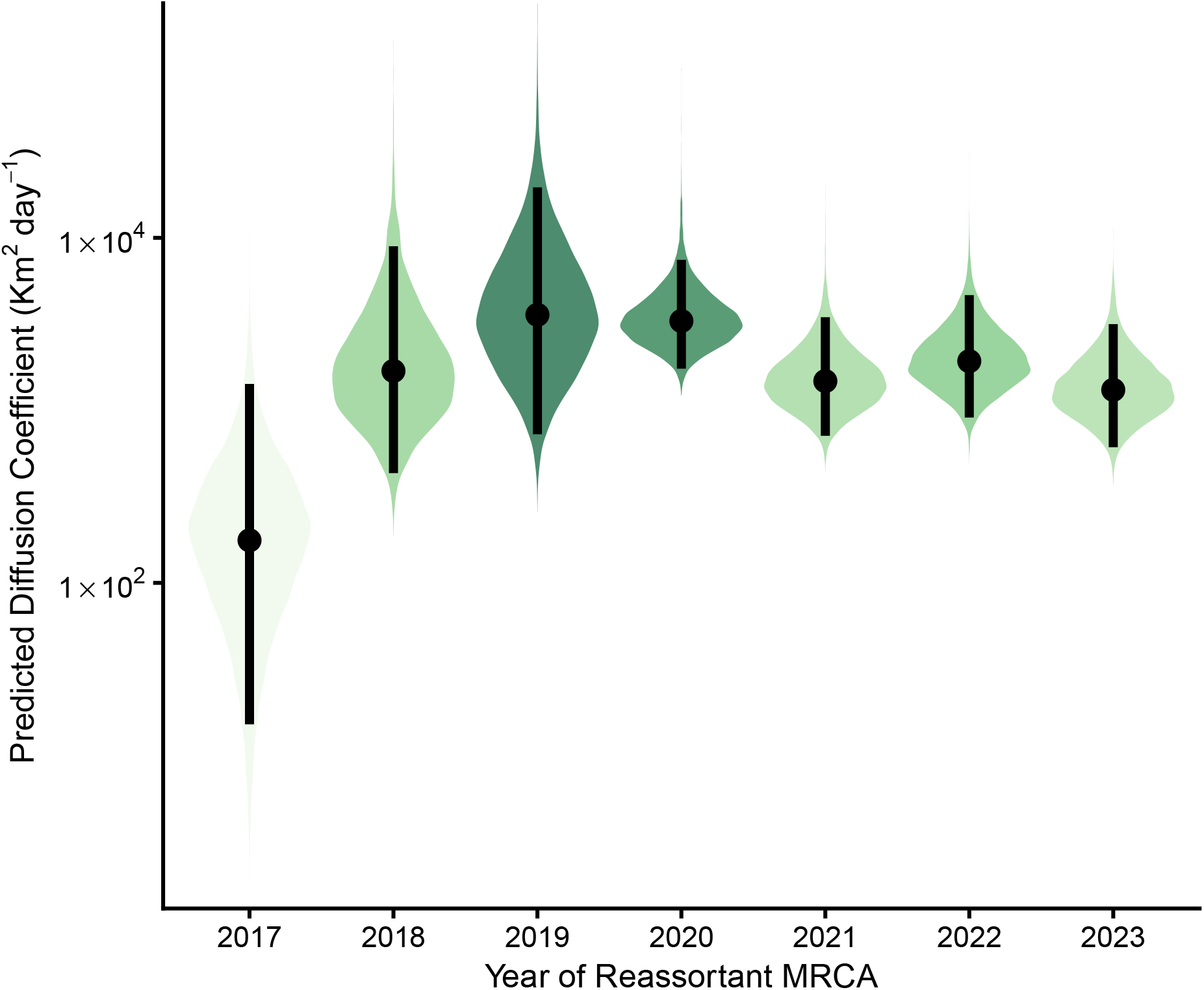
Expectation of the posterior predictive Weighted Diffusion Coefficient distribution, stratified by year of reassortant emergence. The centre dot is the median and the bars correspond to 95% highest posterior density region

**Fig. S21.**
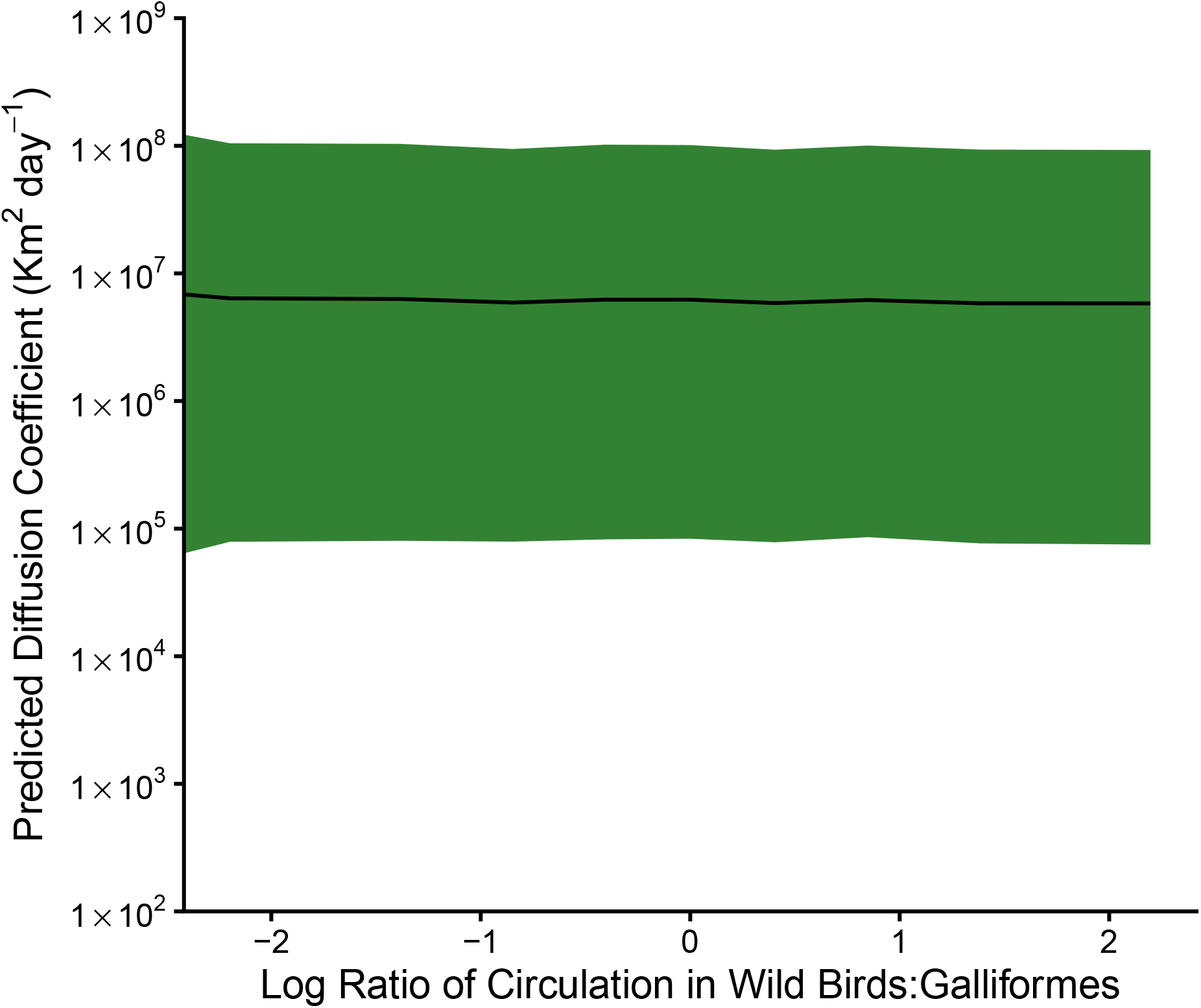
Posterior prediction of the weighted diffusion coefficient as a function of the log ratio of evolutionary time in wild birds to domestic birds. We held persistence at 1 year and marginalised over the remaining variables. The black line is the median of the posterior distribution and shaded region corresponds to 95% highest posterior density.

**Fig. S22.**
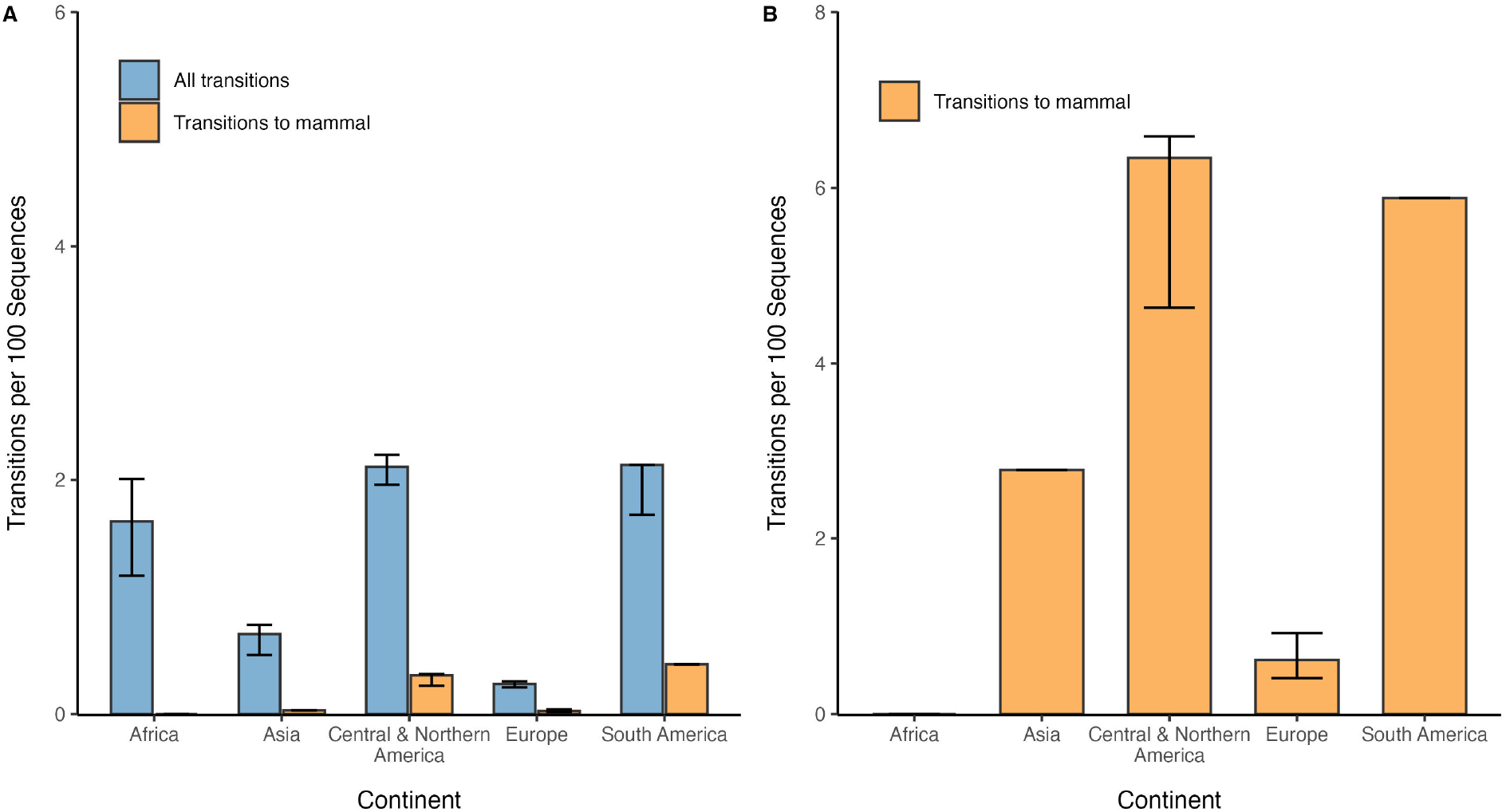
Scaled frequencies of host class transitions for H5 reassortants, stratified by continent. Each plot shows the number of observed host-state transitions for each reassortant, scaled by (A) the total number of sequences and (B) the number of sequences from mammals only. Since the exact number of state-transitions can vary according to each segment, we report the median number of state-transitions for each reassortant and interquartile ranges.

**Fig. S23.**
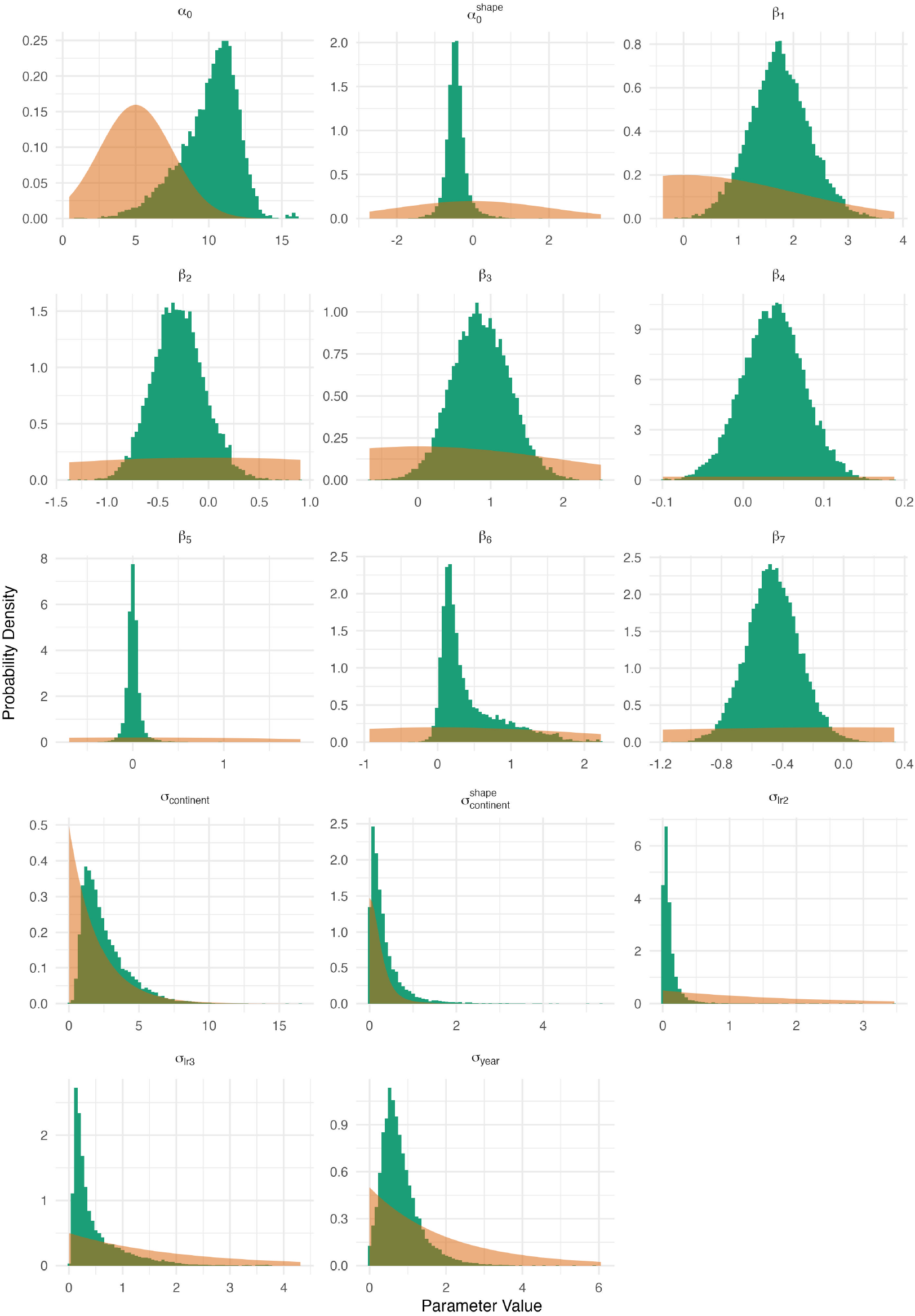
Prior (orange) and posterior distributions (green) for select parameters of the diffusion coefficient model. In all cases the two distributions show minimal overlap, which suggests the posterior distributions are well-informed by the model likelihood.

**Fig. S24.**
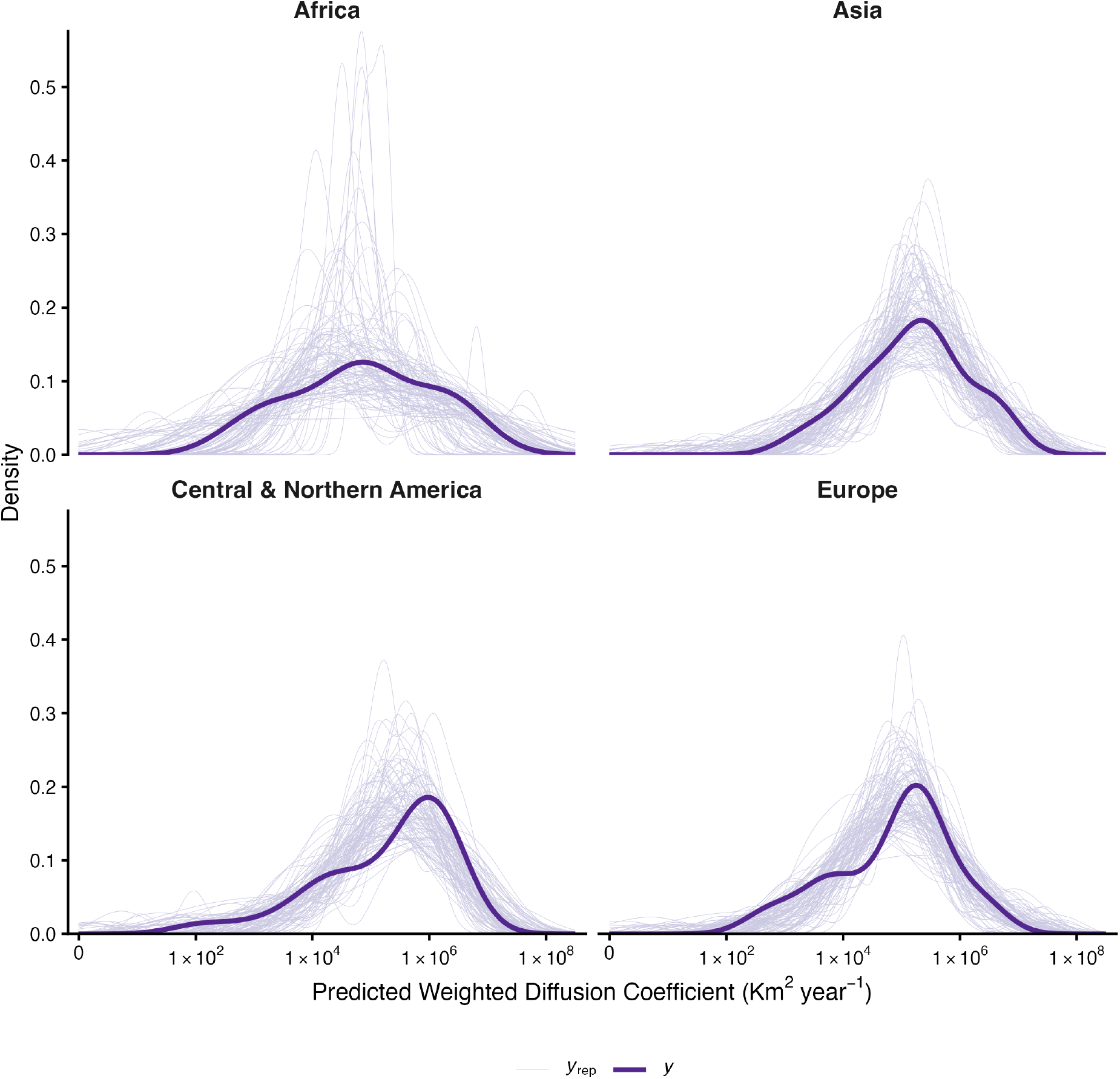
Posterior predictive check for the diffusion coefficient model. Stratified by continent, each thick purple line represents the empirical data distribution. Each thine trace line corresponds to a posterior predictive density, extracted from a single draw.

**Fig. S25.**
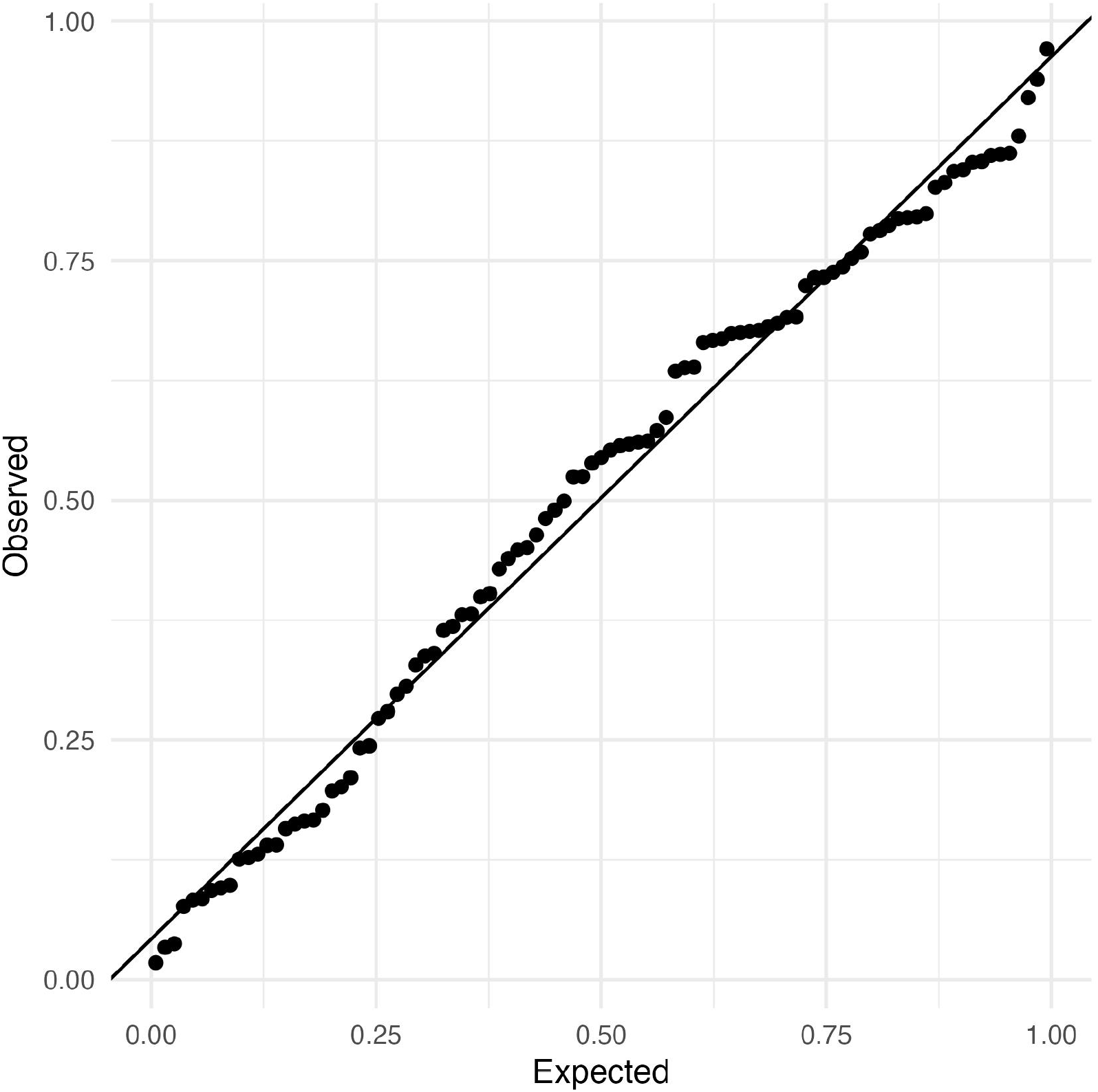
Quantile-Quantile residual diagnostic plot for the diffusion coefficient model. Scaled residuals were calculated using DHARMa [118]

**Fig. S26.**
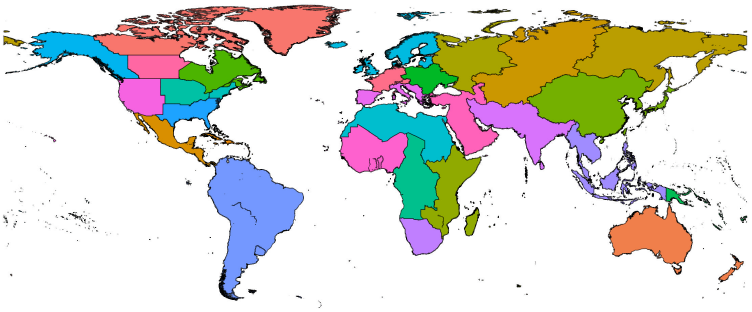
Regions used in the PGLM analysis. We adapted UN geoscheme regions to partition the landmass of the globe into computationally tractable regions.

**Fig. S27.**
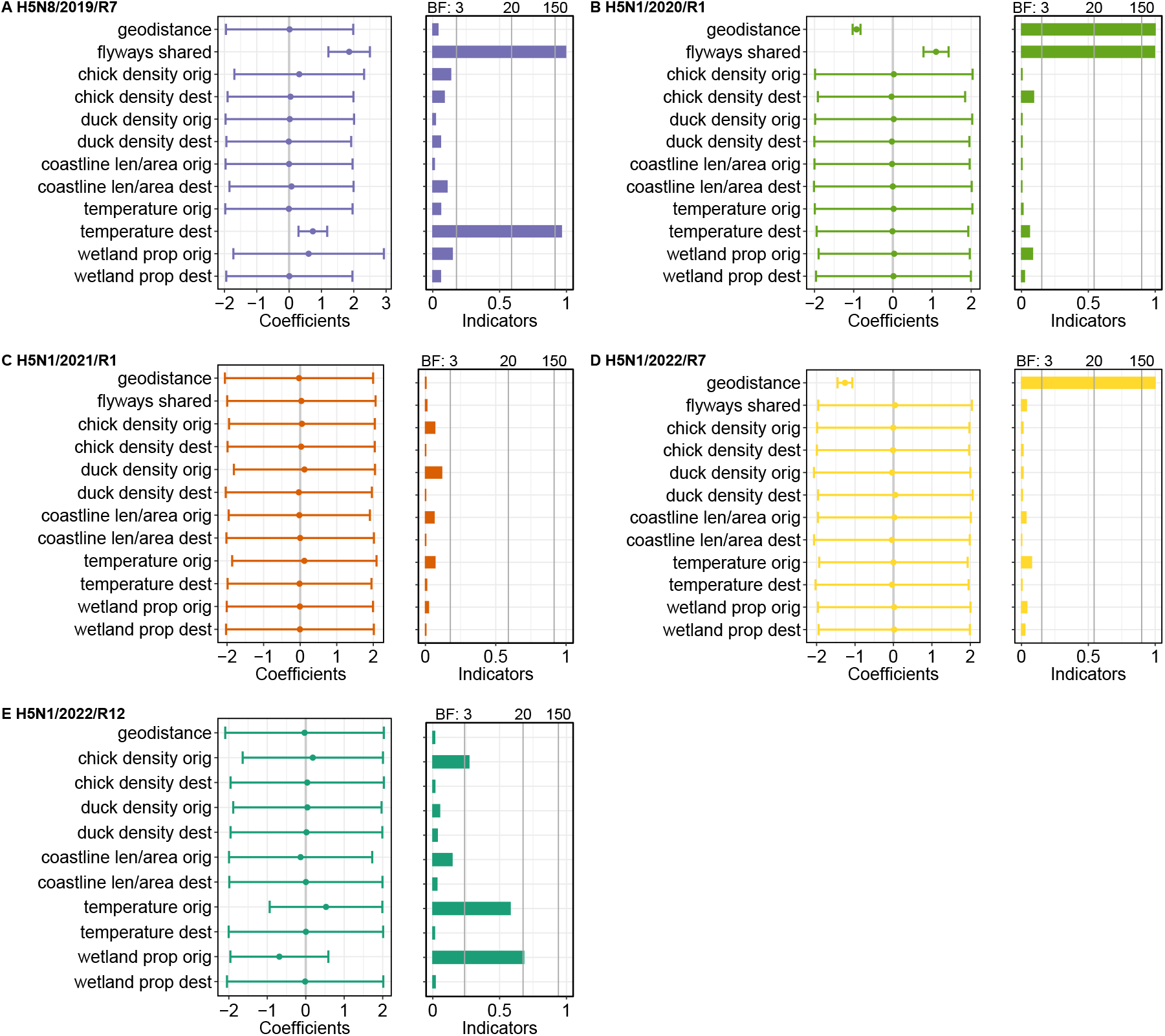
Contributions of predictors for each of five dominant reassortants of H5Nx AIV spread between regions. The virus dispersal is inferred using HA genes by PGLM-extended Bayesian discrete phylogeographic inferences. Highly correlated predictors (absolute Pearson correlation coefficient > 0.7) were analysed in separate PGLMs and the results were aggregated. The predictor related to the number of flyways shared between regions was not included in the PGLM analysis for reassortant H5N1/2022/R12 because the values are identical across regions. The coefficients (left panel) represent the mean size estimate (on a log scale) of the contribution of the predictors with credible intervals. The indicators (right panel) represent the estimated inclusion probability of the predictors. The Bayes factors (BF; grey solid vertical lines in right panel) represent the statistical support for predictors included in the PGLMs. BF *<*3: no or weak support; 3-20: support; and BF >20: strong support.

**Fig. S28.**
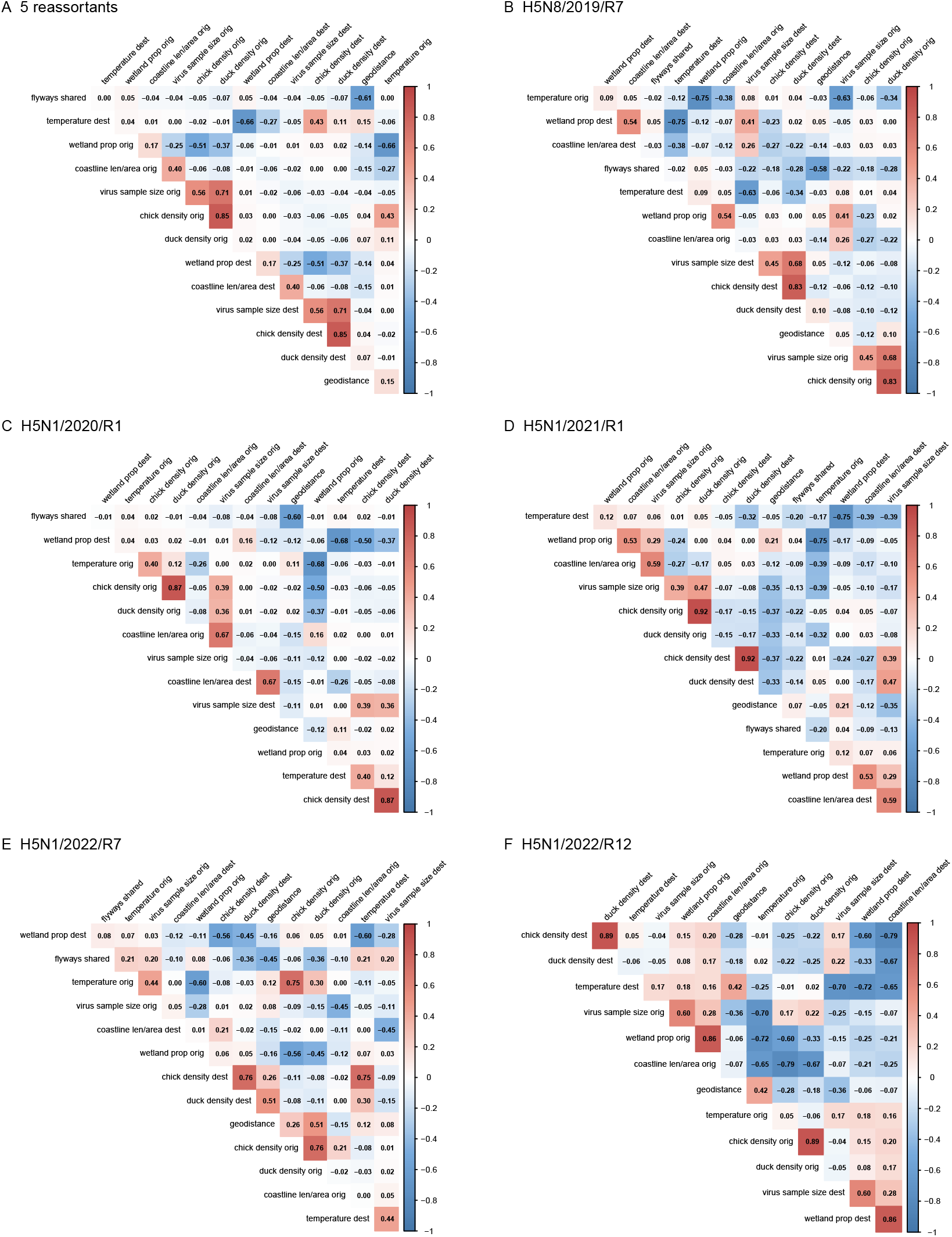
Correlation coefficients of predictors included in the PGLM models used to inform the spread between regions/countries of five major H5 reassortants. The coefficients greater than 0.7 indicate variables that are considered highly correlated. These highly correlated predictors were included in separate PGLM analyses.

**Fig. S29.**
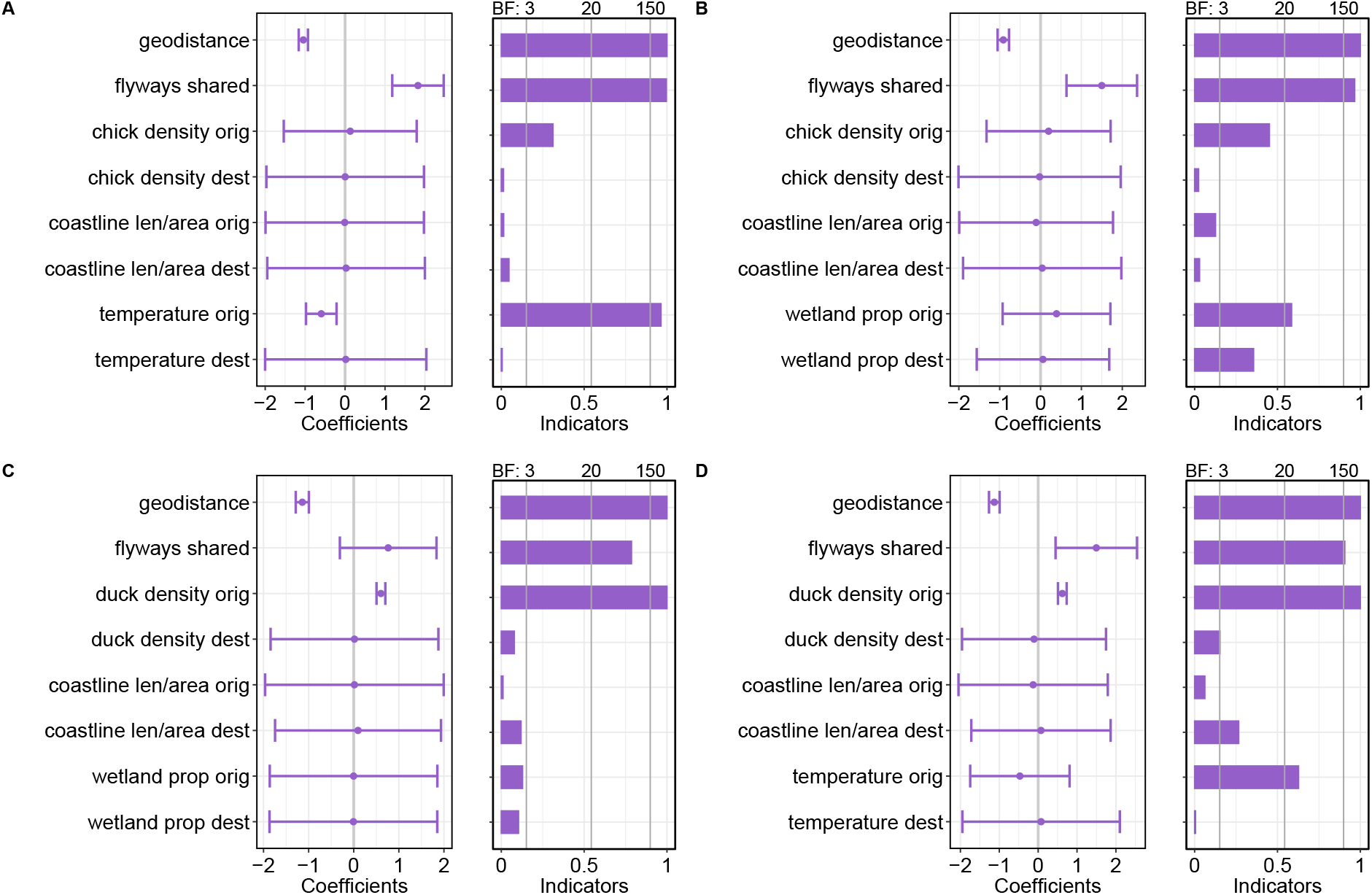
Contributions of predictors for five dominant reassortants of H5Nx AIV spread between regions. The virus dispersal is inferred using HA genes by PGLM-extended Bayesian discrete phylogeographic inferences. Predictors with absolute Pearson correlation coefficient *<* 0.66) were analysed in each PGLM. The coefficients (left panel) represent the mean size estimate (on a log scale) of the contribution of the predictors with credible intervals. The indicators (right panel) represent the estimated inclusion probability of the predictors. The Bayes factors (BF; grey solid vertical lines in right panel) represent the statistical support for predictors included in the PGLMs. BF *<*3: no or weak support; 3-20: support; and BF >20: strong support.

